# Paired oral clinical specimens reveal the underlying ecology supporting the emergence of inflammophilic microbiome communities

**DOI:** 10.64898/2026.02.20.706901

**Authors:** Madeline Krieger, Kristopher A. Kerns, Elizabeth A. Palmer, Jeffery S. McLean, Jens Kreth, Galip Gürkan Yardimci, Justin L. Merritt

## Abstract

**Background:** Inflammatory oral diseases are associated with reproducible shifts from commensal-dominated microbiota toward pathobiont-enriched communities, yet the ecological mechanisms underlying the emergence of inflammophiles remain poorly understood. This study aims to investigate if host-derived inflammatory environments act as selective pressures that restructure microbial metabolism and community organization during disease progression.

**Methods:** We performed 16S rRNA gene sequencing of patient-matched pediatric dental plaque and odontogenic abscess specimens to capture microbial community transitions across an inflammatory ecological gradient. Community ecology modeling and inferred metagenomic analyses were used to identify taxa and functional programs associated with commensal and inflammophilic states.

**Results:** Patient-matched comparisons revealed a reproducible ecological selection gradient linking inflammatory environments to expansion of metabolically specialized inflammophiles and depletion of carbohydrate-utilizing commensals. Commensal-dominated plaque communities exhibited anabolic, carbohydrate-centered metabolic capacity, whereas abscess microbiota were enriched for catabolic metabolism, amino acid fermentation, and antimicrobial resistance, consistent with adaptation to inflammation-driven nutrient landscapes and immune pressure.

**Conclusions:** These findings support a model in which host inflammation drives ecological restructuring of the oral microbiome toward metabolically adapted inflammophilic communities. Defining the metabolic requirements and selective pressures governing these transitions provides a framework for microbiome-directed therapeutic strategies aimed at restoring ecological stability during inflammatory dysbiosis.

## Introduction

The oral cavity supports a complex microbial ecosystem that is second only to the gastrointestinal tract in measures of biodiversity (1). Its varied anatomical landscapes of disparate tooth enamel and epithelial surfaces create distinct ecological niches for oral microbiome communities (2). Among these, oral biofilms (i.e., dental plaque) and odontogenic abscesses develop two discreet community compositions. Dental plaque is primarily composed of commensal species during eubiosis as well as acidogenic and/or proteolytic species during diseased, dysbiotic conditions (2–7). Odontogenic abscess communities are initially seeded by the dental plaque microbiota (8,9) but undergo a dramatic compositional transformation within the abscess environment. This transition is thought to be a consequence of host inflammatory selective pressure (10–12), resulting in communities that become increasingly dominated by a variety of “inflammophilic” microbiota, such as species of *Fusobacterium, Prevotella, Parvimonas* and others (13–18). While such organisms are commonly detected in a variety of studies (5,14,19–21), variability in these results has challenged the discernment of the broader ecological characteristics of inflammophilic microbiomes. Studies of oral inflammatory dysbiosis frequently compare clinical specimens of health vs. disease using patient material derived from separate individuals rather than intra-individual comparisons. This type of sampling can complicate efforts to determine the relationship between host inflammatory responses and microbiome ecology due to the myriad of variables that contribute to community-level fluctuations in the microbiomes of different individuals, such as diet, lifestyle, genetics, and environment (22–25). In response, we collected a cohort of paired disease-free dental plaque and odontogenic abscess clinical specimens concurrently sampled from single individuals to mitigate many of the key limitations of microbiome variability. Additionally, the sequestration and inherent physical separation of abscess and dental plaque communities further alleviates the confounding influence of clinical specimen cross contamination during sampling procedures. From this cohort, we could establish a patient-specific baseline community composition in each dental plaque specimen to examine how its community ecology has evolved within the abscess due to inflammatory selective pressure from the host.

## Results

### 1. Dental plaque and abscess community compositions are distinct

Our cohort of patient-matched dental plaque and abscess clinical specimens was used to directly investigate the ecological influence of host inflammatory pressure on the oral microbiota. Odontogenic abscesses are directly seeded by extant dental plaque communities (8,9) that become sequestered and isolated from external influences once the host assembles walled abscesses far below the gum line (**Fig. 1**). This unique feature of tooth abscess physiology also limits the common problem of specimen cross-contamination with the oral microbiota during clinical sampling, as abscess communities are only revealed following tooth extraction and remain physically separated from surrounding dental plaque throughout the procedure. To investigate the differences between dental plaque and odontogenic abscess communities, we performed 16S rRNA V3V4 gene sequencing of 25 paired, patient-matched plaque and abscess specimens generated from regularly scheduled clinical interventions at the Oregon Health and Science University pediatric dental clinic.

**Figure 1.**
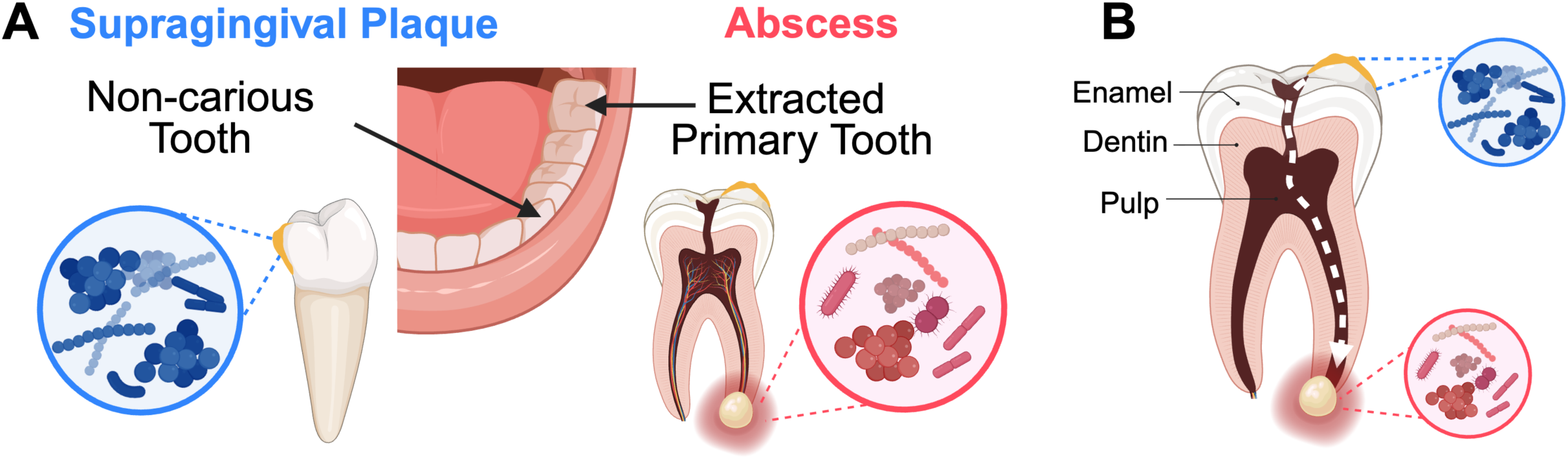
Patient sampling and the ecology of oral abscess formation. (A) The cohort was constructed from patient-matched specimens taken from the supragingival plaque of non-carious primary canines and the abscess material of an extracted primary molar. (B) Caries development begins with demineralization of the enamel surface. If left untreated, the lesion can progress into a deep caries lesion that approaches the tooth pulp chamber. Once the caries lesion has breached the pulp chamber, a direct pathway is created between the oral cavity and the normally sterile tooth pulp, allowing oral microbiota from dental plaque to invade through the caries lesion. Colonization of the pulp chamber initiates an infection that can spread through the root canal to the tooth apex or root furcation, resulting in apical or furcal (pictured) abscess formation.

To assess between-sample variation, we profiled beta diversity using Principal Coordinate Analysis (PCoA) (**Fig. 2A**). The first ordination axis captured true biological variation rather than sequencing artifacts, supported by a minimal correlation with library size (Spearman ρ = 0.18 at the phylum level and –0.25 at the genus level). For comparison, perfect positive or negative correlations are represented by ρ = 1 or ρ = –1, respectively. We further measured the median silhouette width for each community type to assess the ability of PCoA to distinguish plaque and abscess communities. We refer to this metric as a silhouette score. A silhouette score of 1 indicates perfect separation, while a score of −1 indicates no separation. Phylum level analyses yielded silhouette scores of 0.2 for plaque and 0.1 for abscess, whereas genus level analyses yielded scores of 0.38 and 0.28 for plaque and abscess, respectively. Silhouette scores further increased with species-level resolution (0.33 and 0.39 for abscess and plaque communities, respectively) (**Fig. S1A**).

**Figure 2.**
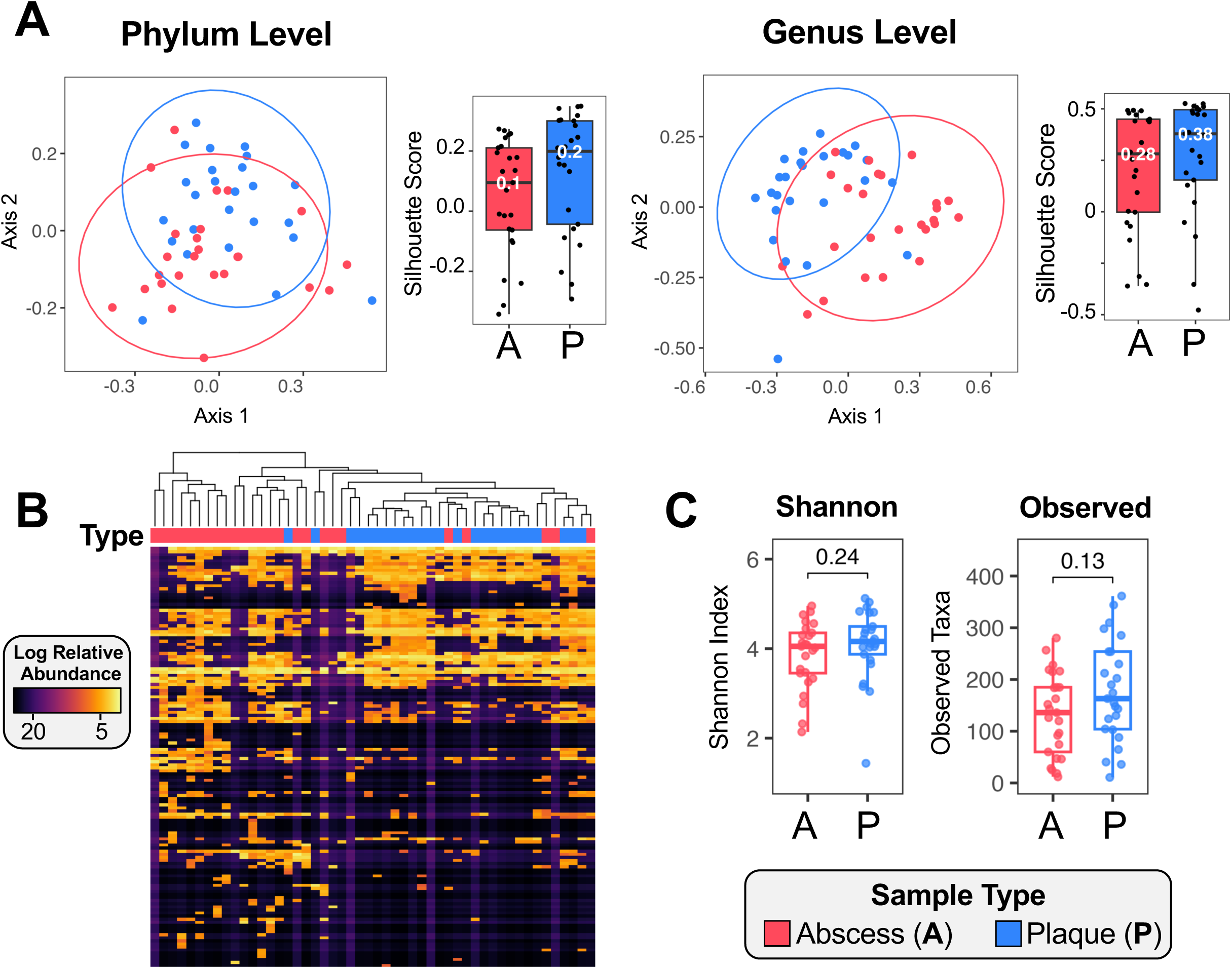
Ecological composition of plaque and abscess communities. (A) Principal coordinate analysis (PCoA) of relative abundance values using Bray–Curtis distance at the phylum (left) and genus (right) levels. Silhouette scores are shown alongside each PCoA plot. (B) Hierarchical clustering (Pearson correlation) of log-transformed relative abundances of genera separates plaque and abscess communities with some overlap. (C) Alpha diversity measured by Shannon and Observed richness indices, with p-values from Wilcoxon Signed-rank tests are displayed above the two groups.

Results of PCoA analysis (**Fig. 2A; Fig. S1A**) indicate that community composition can distinguish between plaque and abscess specimens. Similarly, hierarchical clustering of genus-level relative abundances using Pearson correlation also revealed clear distinctions between plaque and abscess communities (**Fig. 2B**). The same method could distinguish between the communities at species-level resolution as well, albeit with greater variability (**Fig. S1B**). We next compared within-sample diversity using Shannon and Observed richness metrics of the amplicon sequence variants (ASVs) present in each sample (**Fig. 2C**). Neither metric was significantly different between the two communities, suggesting that they are similar in both the total number of unique taxa as well as the evenness of taxa composition in the specimens. We noted a weak trend towards greater biodiversity in the plaque specimens, but it did not reach statistical significance (observed alpha diversity, Wilcoxon Signed-rank p = 0.13) (**Fig. 2C**, right).

### 2. Comparison of taxa enrichment using multiple differential abundance metrics

Given the distinct community compositions found between plaque and abscess specimens, we were next interested in comparing the differential abundances of the specific taxa in the two communities. The overall composition of each community is shown in Figures 3A and 3B. At the phylum level, plaque and abscess communities are broadly similar (**Fig. 3A**). However, at the genus level, compositional distinctions become clearly discernable (**Fig. 3B**). The top 15 genera with the largest significant differences between median values of the two groups were determined using a Wilcoxon Signed-rank test (Benjamini-Hochberg false discovery rate corrected p-value < 0.05) (**Fig. 3C**; **Table S1**), which is straightforward statistical assessment useful for identifying robust, consistent changes across samples. To further confirm that the observed trends were not attributable to major outliers, we also tallied the number of specimens exhibiting enrichment of each genus in the plaque and abscess specimens. These counts are presented as a bubble plot together with the respective enrichment results (**Fig. 3C; Table S2**). For example, genera mainly associated with oral health, such as *Neisseria*, *Lautropia*, *Actinomyces,* and *Rothia* (26), were strongly associated with the plaque specimens, whereas the *Prevotella, Treponema,* and *Dialister* (27,28) genera stratified with the abscesses, in agreement with the prominent inflammophilic species they contain (29,30). We also further resolved these data to the species level (**Fig. 3D; Table S3**). The common health-associated commensal species *Streptococcus oralis* subspecies *dentisani, Neisseria flava, Streptococcus sanguinis* (31), *Lautropia mirabilis,* and *Haemophilus parainfluenzae* (32) all dominated the plaque specimens. Conversely, the oral pathobionts *Prevotella oris*, *Dialister invisus,* and *Alloprevotella tannerae* (29) were strongly associated with the abscesses.

**Figure 3.**
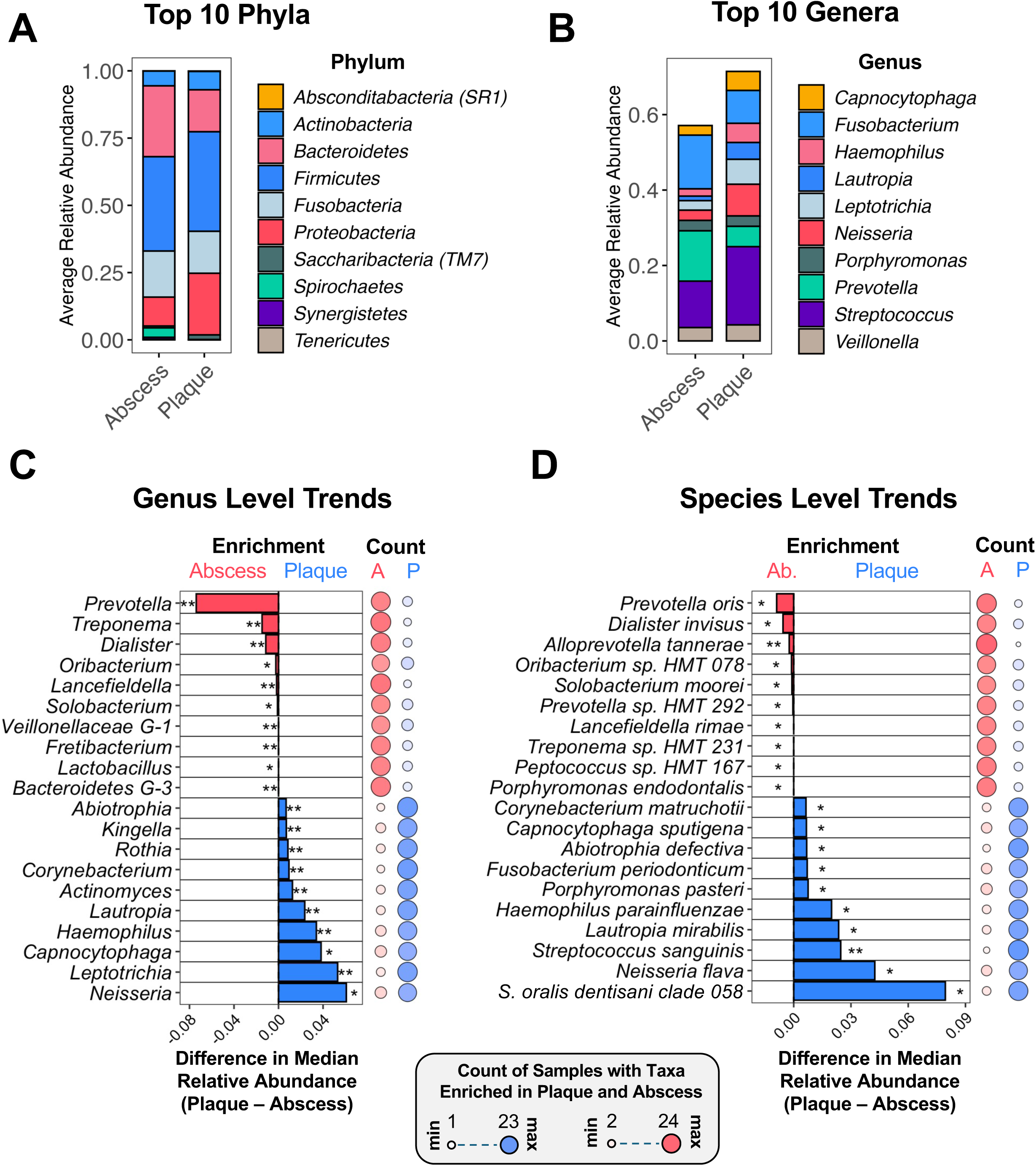
The distribution of taxa within oral plaque and abscess specimens. Average relative abundances of the (A) top 10 phyla and (B) top 10 genera in plaque and abscess specimens. (C) Genus- and (D) species-level enrichment trends. The bar plot (left) is ranked by taxa with the greatest differences between plaque and abscess abundance. The bubble plot (right) signifies the number of samples that have higher relative abundance values of each taxon in the abscess (red) or plaque (blue) communities.

Given the significant differences observed from the Wilcoxon Signed-rank test, we next analyzed the sequence results using differential abundance analytic approaches specifically tailored for microbiome biomarker discovery. Due to the disparate results commonly reported from studies using different differential abundance analysis methods (33,34), we analyzed the plaque and abscess sequence data using three commonly employed approaches: Microbiome Multivariable Associations with Linear Models (MaAsLin2) (35), Analysis of Compositions of Microbiomes with Bias Correction (ANCOMBC2) (36), and Analysis Of Differential Abundance Taking Sample and Scale Variation Into Account (ALDEx2) (37). MaAsLin2 and ALDEx2 uses generalized linear models, with MaAsLin2 allowing incorporation of metadata covariates. ANCOMBC2 uses a mixed-effects framework with covariate consideration. Applying each of the differential abundance algorithms resulted in slightly different outcomes: MaAsLin2 identified 4 phyla, 30 genera, and 54 species, ANCOMBC2 yielded 2 phyla, 11 genera, and 20 species, and ALDEx2 identified 4 phyla, 25 genera, and 44 species (**Table S4**). Taxa identified by all three methods were labeled as high confidence, medium confidence taxa were identified by two methods, and low confidence taxa were unique to a single algorithm. At the phylum level, we identified 2 high-confidence taxa: *Proteobacteria* (plaque enriched) and *Spirochetes* (abscess enriched) (**Fig. 4A**), along with one medium confidence abscess-enriched phylum, *Synergistetes.* Low confidence enrichment of *Tenericutes* and *Saccharibacteria* were detected in the plaque and abscess communities, respectively. At the genus level, 6 high confidence taxa were identified: *Leptotrichia*, *Corynebacterium*, and *Actinomyces* for dental plaque specimens and *Veillonellaceae G-1*, *Dialister*, and *Treponema* for abscesses (**Fig. 4B**). 13 medium confidence enriched taxa were found in the plaque compared to 2 in the abscess (**Fig. 4B**). As in Figures 3C and 3D, a count of samples with enrichment of each taxon in the specimens is shown to the right of each plot, with specific count values presented in Table S5. Dot plots for the relative abundance/total sum scaling (TSS) values of these high and medium confidence genera indicate the results were not driven by outliers in the data (**Fig. S2A; Fig. S2B**).

**Figure 4.**
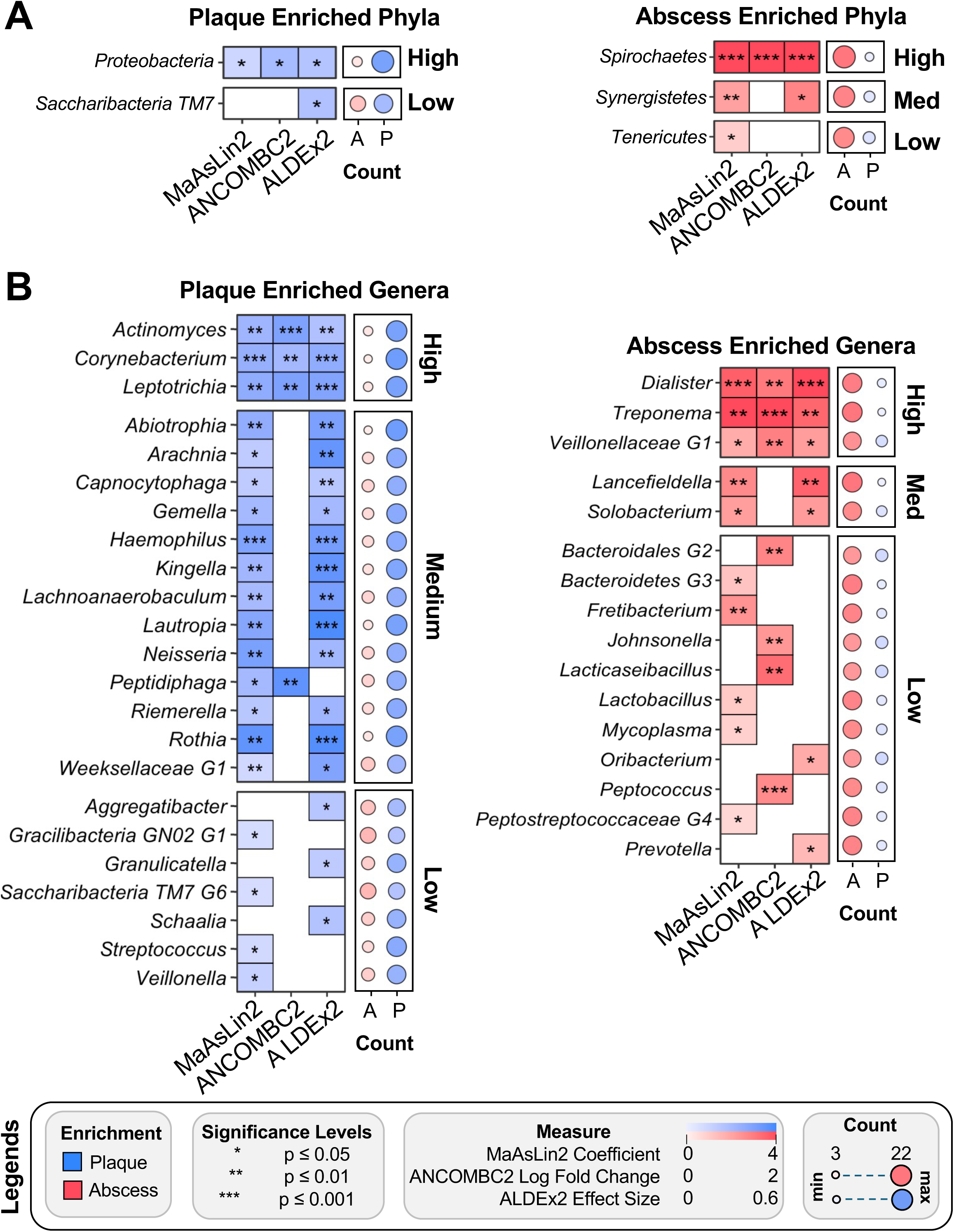
Differentially abundant phylum- and genus-level taxa in plaque and abscess communities. Results from three differential abundance methods (MaAsLin2, ALDEx2, ANCOMBC2). Significant enrichment of a taxon via all three methods resulted in its classification as high confidence, significant enrichment in two methods resulted in a medium confidence classification, and significance from a single method was classified as low confidence. The bubble plot to the right of each plot signifies the number of samples exhibiting enrichment in the abscess (red) or plaque (blue) communities. (A) Phylum-level results. (B) Genus-level results.

When examining species-level resolution (**Fig. 5**), high confidence enrichment of the oral commensal species *Streptococcus sanguinis*, *Rothia aeria*, *Corynebacterium durum,* and *Porphyromonas pasteri* (38) were all detected in the plaque specimens as well as the uncharacterized organism *Alloprevotella* sp. HMT 473. Interestingly, the unclassified taxon *Oribacterium HMT 078* is the only organism found to be enriched in the abscess by all three differential abundance analysis approaches. It is worth noting that we detected a weak enrichment of *Fusobacterium nucleatum* in the abscess (**Fig. 5**) that was not captured by the genus-level differential abundance analysis methods (**Fig. 4**), despite known associations of *F. nucleatum* with oral inflammatory diseases (39). Intrigued, we further confirmed these results using TSS and center log ratio (CLR) transformations to visualize the distribution of fusobacteria within the plaque and abscess specimens (**Fig. S3**). A recent subspecies-level analysis of *F. nucleatum* in oral clinical specimens identified exceptionally biased subspecies-level stratification of *F. nucleatum* within patient-matched health vs. disease specimens (40). Thus, *F. nucleatum* sequence data would likely require further resolution to the subspecies level to discern plaque- and abscess-specific enrichment trends. However, such resolution is poorly captured by 16S rRNA gene sequencing (40,41).

**Figure 5.**
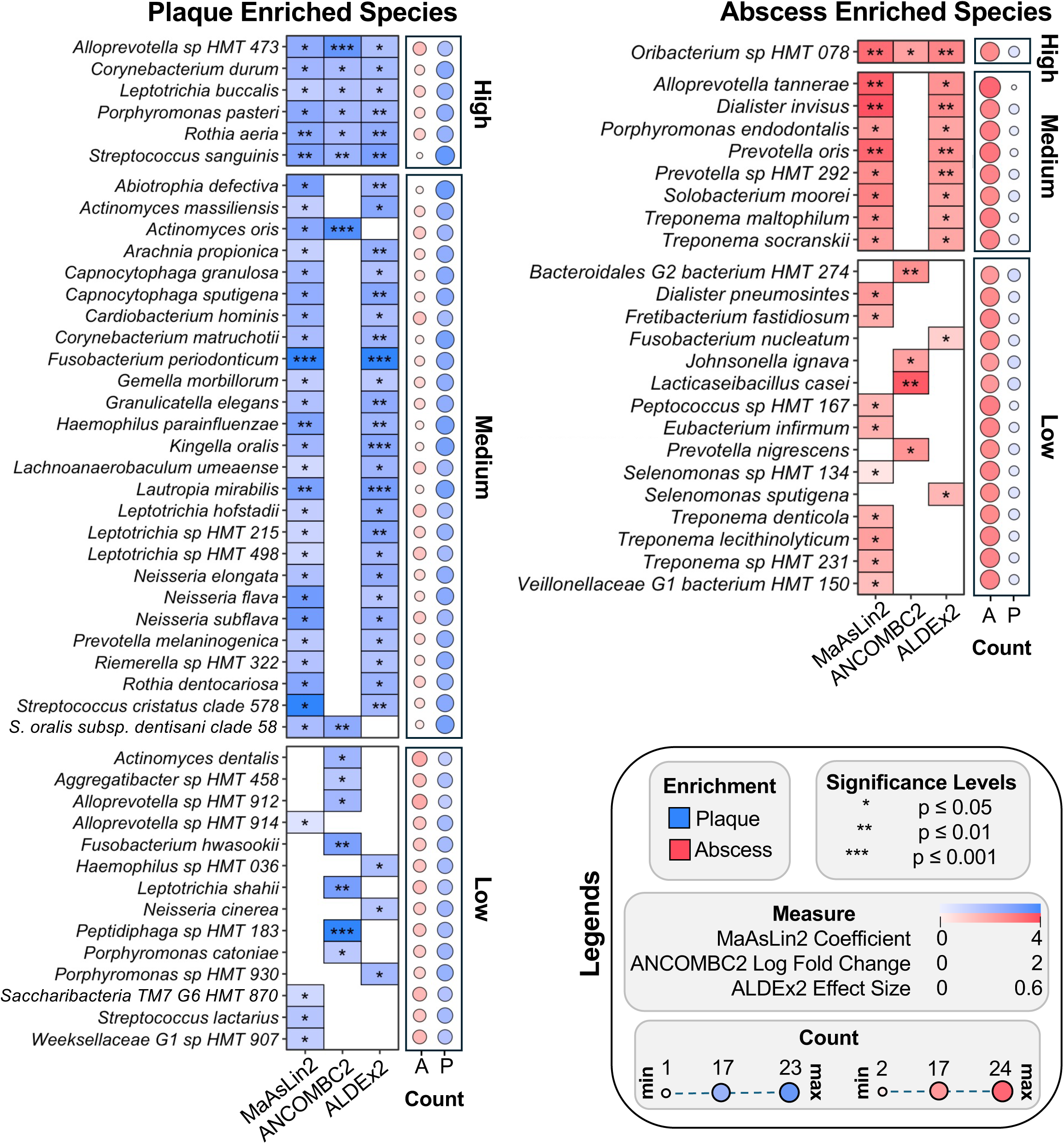
Differentially abundant species in plaque and abscess communities. Results from three differential abundance methods (MaAsLin2, ALDEx2, ANCOMBC2). Significant enrichment of a taxon via all three methods resulted in its classification as high confidence, significant enrichment in two methods resulted in a medium confidence classification, and significance from a single method was classified as low confidence. The bubble plot to the right of each plot signifies the number of samples exhibiting enrichment in the abscess (red) or plaque (blue) communities.

### 3. Topic modeling reveals community-level drivers within plaque and abscess communities

While differential abundance analyses identified specific plaque- and abscess-enriched taxa within the specimens, these analyses do not capture broader ecological trends in the communities, which is a key facet defining eubiosis vs. dysbiosis. Therefore, we subsequently employed two different machine learning approaches that can identify broader community-level trends in the specimens using both supervised and unsupervised methods.

First, we applied random forest classification with k-fold cross-validation. This approach partitions the dataset into multiple training and testing folds, enabling the model to learn classification patterns from the training data followed by an evaluation of the performance of the trained model on the remaining test set. This approach is ideally suited to balance bias variance tradeoffs in cohorts with moderate sample sizes; namely, we can train models on a large training set, while assessing the generalizability of the trained models across different folds. We elected to use fourfold cross validation to train random forest models that can distinguish between clinical specimen types in which ¾ of the sequence data was used for training and the remaining ¼ used for analysis (**Fig. 6A)**. Using genus-level compositional data, our model correctly predicted specimen type with 75-83% accuracy (evaluated by the area under the curve) across the 4-folds (**Fig. 6B**). Although taxa were generally consistent across folds, some variability was observed (**Fig. 6C**). For example, *Dialister* ranked as the strongest predictor in folds 2 and 4, but fifth and seventh in folds 1 and 3, respectively. *Treponema* was the second most important feature in folds 1 and 4, but it ranked 10^th^ and 6^th^ most important in folds 2 and 3, respectively. Fold 2, which had the highest accuracy (83%), prioritized the genera *Dialister*, *Corynebacterium*, and *Lautropia* as the 1^st^–3^rd^ most important specimen type predictors. These genera were also significantly enriched in our prior analysis (**Fig. 3C**; **Fig. 4B**). Interestingly, the 4^th^ strongest predictor was *Lachnoanaerobaculum,* a genus not previously identified in our differential abundance analyses. Although random forest performed well at the genus level, it was slightly less consistent using species-level data, with accuracy ranging from 55-85% (as measured by area under the curve) (**Fig. S4A**). Much of this variability in community composition could be specifically attributed to fold 1. Despite this, *Streptococcus sanguinis* and *Dialister invisus* remained among the top 16 predictors between all four folds (**Fig. S4B**), which is highly consistent with our differential abundance analysis (**Fig. 5**). Based upon the cohort size and large number of features, some variation across different folds was expected, even for broadly consistent predictors.

**Figure 6.**
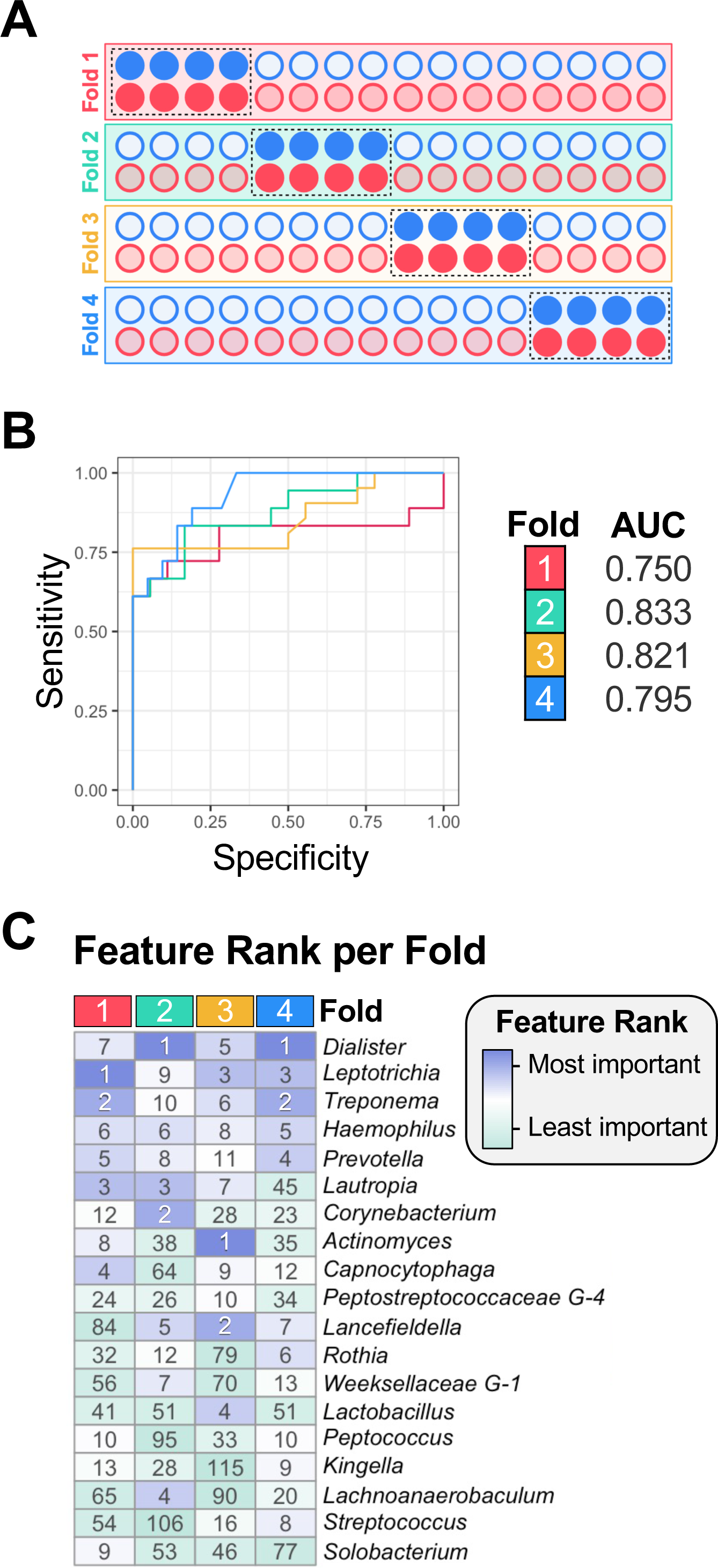
Genus-level community composition predicts clinical specimen type. A random forest classifier was trained on genus-level relative abundances using 4-fold cross validation. (A) A schematic illustrating how 4-fold cross validation was employed to divide the plaque (blue circles) and abscess (red circles) samples into four equal folds, with ¼ of the data (dark colored circles) withheld to test the accuracy of the algorithm trained on the remaining ¾ of the data (light colored circles). (B) Receiver operating characteristic (ROC) curves from 4-fold cross validation demonstrate strong performance (area under the curve, or AUC values of 0.75–0.83). (C) Heatmap of feature importance rankings across folds, with top predictors in purple and lower-ranked features in green. The rank of feature importance is shown inside each cell.

The observed variability in the species-level random forest results led us to pursue a new analytic approach that we predicted would be ideally suited for microbiome datasets: Latent Dirichlet Allocation (LDA) topic modeling. LDA is an algorithm originally developed for natural language processing to organize text-based documents into underlying themes or topics in an unsupervised fashion, with no prior knowledge of the underlying composition and contents of each topic. LDA has been applied to microbiological datasets in a few prior studies (42–46), but it remains an uncommon analytical approach in the field. Our adaptation of this algorithm employs scaled relative abundance values of microbiome taxa and is summarized in Figure 7A. In contrast to standard differential abundance methodologies (**Fig. 4**; **Fig. 5**), LDA can learn polymicrobial signatures of co-occurring taxa within datasets without each member of the signature being present in all specimens, potentially mitigating the impact of clinical specimen variability while learning signatures that can be broadly shared across clinical specimens in the cohort. After stratifying the microbiome into latent community structures (topics), we next examined the degree of sample membership within each topic (gamma). Using goodness of fit metrics (**Fig. S5A**), we created a species-level LDA model with six topics (**Fig. 7B**). A Wilcoxon Signed-rank test yielded a highly significant separation (p < .001) between several plaque-specific (topics 2 and 4) and abscess-specific (topic 3) topics (**Fig. 7C; Fig. S6**). Non-significant topics that capture shared signatures of both communities were also identified (topics 1, 5, and 6) (**Fig. 7C; Fig. S7**). Uniform Manifold Approximation and Projection (UMAP) of gamma values effectively separated plaque and abscess communities with moderate silhouette scores (abscess = 0.25; plaque = 0.5) (**Fig. 7D**). The distribution of relative abundances for the top taxa in each topic is presented in Figure S6, revealing clear enrichment patterns in the plaque- and abscess-dominated topics. The plaque-specific topics are principally driven by the obligately commensal species *Neisseria flava* (topic 2) and *Streptococcus oralis* subspecies *dentisani* (topic 4), whereas *Fusobacterium nucleatum*, *Prevotella nigrescens*, and *Bacteroidales* G2 HMT 274 were the most prominent within the abscess-specific topic 3 (**Fig. 7E**). Interestingly, topic 3 is also far less biased for a single species compared to the plaque-specific topics 2 and 4. The top drivers of the shared signatures (topics 1, 5, and 6) are shown in Figure S7.

**Figure 7.**
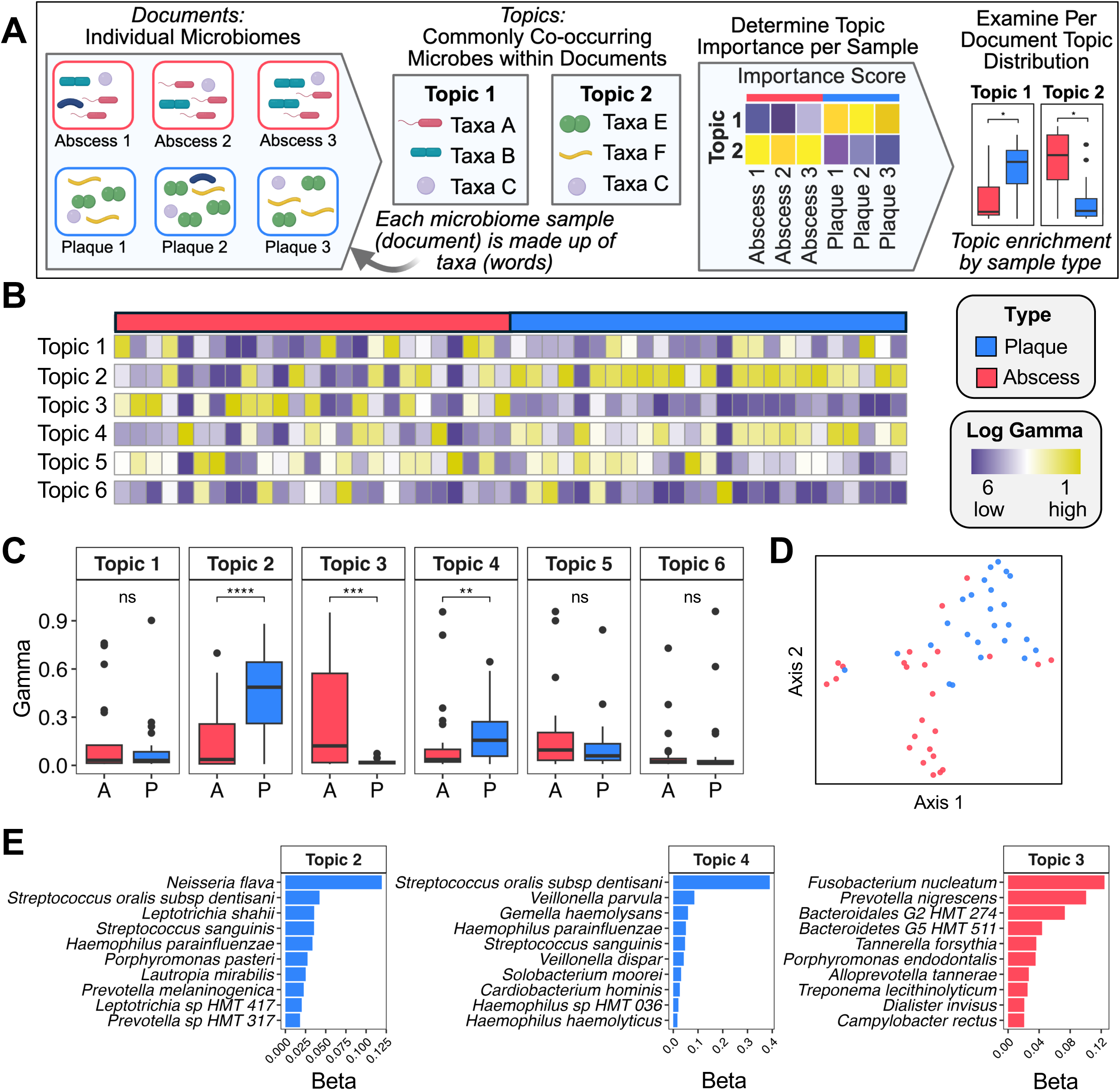
Species-level topic modeling of plaque and abscess specimens. (A) An illustration demonstrating how Latent Dirichlet Allocation (LDA) can be applied to microbiome data to elucidate community-level signatures. (B) Heatmap of gamma scores across topics. (C) Boxplots of sample membership (gamma values), with the results of a Wilcoxon Signed-rank test displayed above (*** = p < .001, * = p < .05). (D) Uniform manifold approximation and projection (UMAP) of gamma scores demonstrate clear plaque–abscess separation. (E) The importance (i.e., beta) scores of the top species driving each topic. Only the results from the topics significantly enriched in plaque communities (topics 2 and 4) and abscess communities (topic 3) are shown. The beta scores of the shared topics are shown in Figure S7.

For comparison, we also repeated the LDA analysis using genus-level data, resulting in three plaque-specific topics (topics 1, 4, and 6), two abscess-specific topics (topics 3 and 5), and two shared topics (topics 2 and 7) (**Fig. S5B; Fig. S8A–C**; **Fig**. **S9**). At this lower level of phylogenetic resolution, LDA analysis still identified key genera driving differential community structures, such as *Neisseria* (plaque) and *Fusobacterium* (abscess) (**Fig. S8C**). However, there was comparatively less overall distinction between the plaque and abscess specimens, presumably due to the mixture of oral eubiont and pathobiont species found in many genera. For example, the genus *Fusobacterium* appeared as a main driver in both the abscess-specific topic 5 and shared signature topic 2 (**Fig. S8; Fig. S9**). This illustrates the flexibility of LDA for microbiome analysis, as the same taxon can appear in multiple topics. Collectively, these results support the utility of topic models for revealing polymicrobial signatures present within complex microbiome communities.

### 4. Biochemical pathway analysis of plaque and abscess communities implicates environmental selection

Inflammophilic communities are thought to have an obligate dependence upon host responses to provide the nutrients required for proliferation and survival (16,47). If true, one would predict that such a relationship would correlate with the metabolic pathways enriched in inflammophilic communities. Consequently, we applied two independent algorithmic approaches to investigate inferred metagenomic profiles from the species compositional data. We first employed PICRUSt2 (48) to profile the distribution of enriched functions listed in the KEGG Orthology (KO) database. PICRUSt2 aligns ASVs onto a reference phylogenetic tree and subsequently infers gene and pathway content based upon closely related genomes. MaAslin2 was used to identify specific enrichment of KOs in the plaque and abscess specimens (**Table S6**). The enriched KOs were next analyzed for their contributions to larger KEGG pathways. This analysis yielded 23 identified KEGG pathways in dental plaque communities compared to only 13 in the abscesses, suggesting a greater metabolic heterogeneity within dental plaque communities (**Fig. 8; Table S7**). In further support of this notion, the genes identified by PICRUSt2 for the dental plaque-enriched pathways are primarily involved in biosynthetic anabolic metabolism (14 anabolic vs. 8 catabolic pathways), whereas the abscess-enriched pathways are extremely biased for catabolism (2 anabolic vs. 11 catabolic pathways). Furthermore, the KOs enriched in dental plaque communities largely reflect the Gram-positive species bias in these specimens (**Table S7**). For example, genes involved in peptidoglycan biosynthesis are enriched as well as the genes used for the biosynthesis of the cell wall lipoglycans of oral *Corynebacterium* species (49–53). Likewise, the genes identified in the quorum sensing category are primarily derived from streptococci (**Table S7**). Dental plaque communities also exhibit a specific enrichment of biosynthetic anabolic pathways for various amino acids, nucleosides, and cofactors. Energy generation in dental plaque communities is heavily biased for glycolysis and oxidative phosphorylation (54) of diverse carbohydrates typically encountered in the diet, including starch, fructose, sucrose, galactose, mannose and others as well as a concomitant enrichment of the phosphoenolpyruvate:sugar phosphotransferase systems responsible for importing these sugars from the environment (55). Overall, the pathway enrichment results indicate that dental plaque communities are largely microaerophilic/facultative and metabolically interdependent, cooperatively synthesizing a large number of the key metabolites required for active growth. In contrast, pathways specifically enriched within abscess communities reflect a considerably different ecology and growth strategy. The exceptional bias for catabolic pathway enrichment suggests that abscess communities exhibit a much greater reliance upon the environment to provide the essential metabolites needed for growth. Unlike dental plaque communities, abscess communities generate energy principally from anaerobic amino acid fermentation, including phenylalanine, tyrosine, and tryptophan (xylene degradation pathway), glutamate (C5-branched dibasic acid pathway; glyoxylate and dicarboxylate pathway), cysteine, methionine, glycine, serine, threonine, arginine, and proline (**Table S7**). Interestingly, the arginine and proline metabolism KEGG category was also similarly enriched in the dental plaque communities (**Fig. 8**). However, the genes listed in this category for the dental plaque microbiota are distinct from those enriched in the abscess communities and are highly likely to be employed for polyamine biosynthesis, rather than ATP generation via arginine and proline catabolism (**Table S7**). Abscess communities also exhibited an enrichment of butanoate metabolism, which is a key ATP-generating pathway used during the fermentation of glutamate, lysine, histidine, cysteine, serine, and methionine (56). Interestingly, the butyrate excreted from butanoate metabolism can directly modulate immune responses and even the epigenetic programming of host cells, potentially influencing host responses to inflammophilic abscess communities (57). Further evidence for abscess community dependence upon host-derived proteins can be found among the identified genes categorized in pentose and glucuronate interconversions as well as fructose and mannose metabolism due to their roles in the catabolism of unusual sugars commonly present within host protein glycosylations, such as fucose (58) (**Fig. 8; Table S7**). Conspicuously, abscess communities also exhibit an enrichment of genes involved in cationic antimicrobial peptide (CAMP) resistance, which presumably provides a specific survival advantage when challenged by innate immunity in inflammatory environments (59–61).

**Figure 8.**
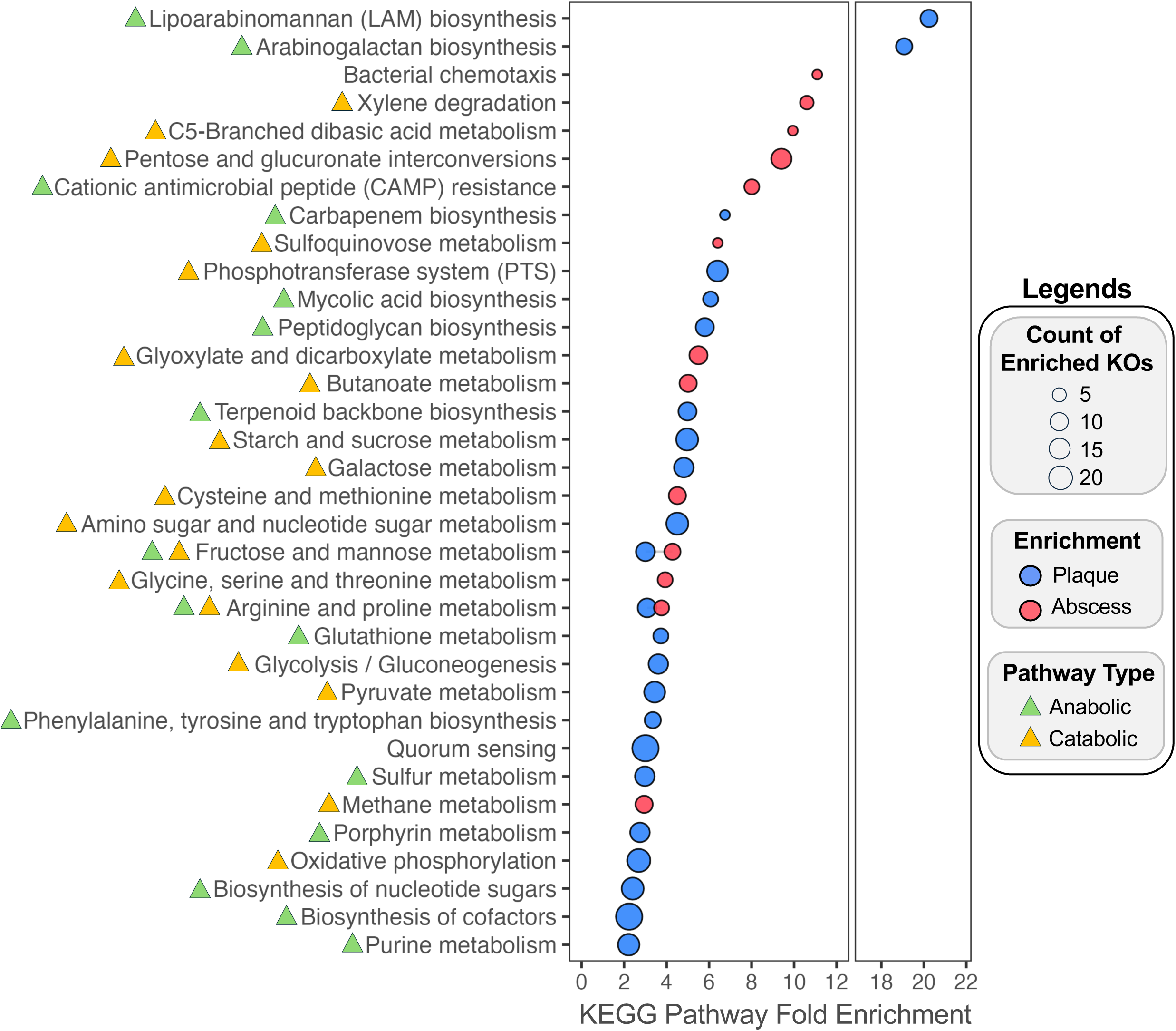
Enrichment of KEGG pathways within plaque and abscess communities. KOs identified by PiCrust2 as differentially enriched in plaque and abscess samples (MaAsLin2 adjusted q value < .05) were used as inputs to identify the associated KEGG pathways. The x-axis displays fold enrichment of the KEGG pathway. Dot size corresponds to the number of unique KOs that were identified as being part of each KEGG pathway (y-axis). Triangle color next to each KEGG pathway category indicates whether the pathway is anabolic (green), catabolic (orange), or contains components of both.

As a separate assessment, we repeated the biochemical pathway analysis using a second approach via the Anvi’o pipeline (62). Publicly available genome sequences of representative enriched species (**Fig. 5**) were downloaded from NCBI RefSeq (**Table S8**). Functional enrichment profiling of a pangenome constructed using Anvi’o (63,64) revealed selective enrichment of multiple KOs (q-value < 0.05) in both the plaque and abscess communities due to the presence or absence of specific gene clusters in each genome (**Table S9**). We applied the same KEGG pathway classification analysis to these plaque- and abscess-specific KOs (**Table S10**). The results largely agree with those of the PICRUSt2 analysis, primarily for the dental plaque communities, whereas far fewer abscess-enriched pathways were detected (**Fig. S10**). However, PICRUSt2 and Anvi’o analyses of abscess communities both similarly detected an enrichment of genes related to motility (i.e., chemotaxis and flagellar assembly) as well as glycine degradation (glycine, serine, and threonine metabolism and one carbon pool by folate) (**Fig. 8; Table S7; Table S10**). The Anvi’o analysis identified two pathways that seemingly contradicted the PICRUSt2 results (C5-branched dibasic acid metabolism and terpenoid biosynthesis) (**Fig. 8; Fig. S10**). However, none of the genes enriched in either category are shared between the PICRUSt2 and Anvi’o outputs (**Table S7; Table S10**). Furthermore, the abscess-enriched genes listed for C5-branched dibasic acid metabolism primarily function in amino acid degradation, whereas those identified in the plaque communities are related to amino acid biosynthesis (**Fig. 8; Fig. S10**). Enrichment of terpenoid biosynthesis genes in plaque communities is primarily attributable to the mevalonate pathway, whereas in the abscess, the identified genes largely function in the non-mevalonate pathway (**Table S7; Table S10**). Thus, no obvious contradictions were detected between the PICRUSt2 and Anvi’o analyses.

## Discussion

In this study, we present a comprehensive ecological analysis of microbiome communities present within a unique cohort of patient-matched clinical specimens of disease-free dental plaque and odontogenic abscess. Prior studies have demonstrated that human dental plaque is dominated by fast-growing taxa like *Streptococcus* and *Actinomyces* (20,65–68) and odontogenic abscess communities are mostly comprised of fastidious, slower growing inflammophilic taxa (17,69). A unique aspect of the odontogenic abscess is that the oral microbiota seeding the infection become shielded from outside influence following their sequestration within the abscess, whereby they subsequently develop under the exclusive influence of the selective pressures within the inflammatory abscess environment (**Fig. 1**). This provides a well-controlled experimental platform to observe how inflammatory pressure from the host remodels a eubiotic microbiome community into a dysbiotic and inflammophilic state. Variability among the results from extant oral health vs. inflammatory disease microbiome data is a common challenge that obscures our understanding of the ecological connection between host inflammatory responses and the emergence of inflammophilic microbiome communities. By leveraging a cohort of patient-matched dental plaque and odontogenic abscess specimens, we could directly observe this remodeling of community ecology while minimizing the confounding influences of inter-individual microbiome variability and specimen contamination during clinical sampling procedures.

Using multiple independent computational techniques, we demonstrated that community composition reliably stratifies plaque vs. abscess microbiomes. Our results also highlight the limitations encountered when comparing the results from studies employing disparate differential abundance analysis methods (33,34). We applied three widely used algorithms to the same sequence data (MaAsLin2, ALDEx2, and ANCOMBC2) and obtained a number of both overlapping and divergent results. Of the three approaches, MaAsLin2 yielded the highest level of significance in the results across almost all confidence levels, even when applying a more stringent significance filter (q-value <.05) than the default (q-value <.25). The only exception was in the low confidence plaque-enriched species, where ANCOMBC2 identified a greater number of significantly enriched taxa than MaAsLin2 (**Fig. 5**). The ideal differential abundance algorithm balances sensitivity and specificity to avoid the over-identification of spurious taxa and the under-identification of taxa with meaningful biological variation. By combining three independent algorithms (33), it was possible to mitigate these limitations while also revealing the discrepancies between them.

Due to the limited species resolution of 16S rRNA V3V4 gene sequence data for certain bacterial species, some species-level trends may have been poorly captured. Despite this, we identified multiple high confidence commensal species in dental plaque (**Fig. 5)**, such as *Streptococcus sanguinis* (31), *Rothia aeria* (70), and *Corynebacterium durum* (71). Conversely, only the unnamed oral taxon *Oribacterium* HMT 078 met our most stringent criteria for high-confidence abscess enrichment. The *Oribacterium* genus is poorly characterized overall, but a variety of *Oribacterium* species are commonly detected in microbiome studies, often associating with mucosal inflammation (72–79). While little is known about *Oribacterium* HMT 078, it is worth noting that this organism was recently reported to be highly enriched in severe early childhood caries lesions together with *Lacticaseibacillus casei* (80). Such deep caries lesions typically precede most endodontic tooth abscesses (**Fig. 1**), suggesting a potential explanation for the observed enrichment of *Oribacterium* HMT 078 in the abscess specimens. For example, the *Lactobacillus* and *Lacticaseibacillus* genera exhibited weak but detectable enrichment within odontogenic abscesses (**Fig. 3C**; **Fig. 4B**), even though neither genus is known to contain oral inflammophiles. Due to their highly acidogenic/aciduric nature, a variety of *Lactobacillus* and *Lacticaseibacillus* species commonly accumulate to abnormally high levels within deep caries lesions, placing an abundance of these species in close proximity to developing abscess communities (80,81) (**Fig. 1**). We propose that a similar phenomenon may be at least partially attributable for the enrichment of *Oribacterium* HMT 078 in the abscess specimens.

Given the limitations of differential abundance analyses for diverse microbiome communities, we adapted an LDA topic modeling approach to identify multi-taxa signatures from relative abundance data. A key strength of LDA analysis is that it is far more permissive of the natural variability that exists between human microbiomes (**Fig. 7A**). This flexibility was especially apparent when applied to genus-level data. For example, the plaque-specific topic 1 was largely driven by *Leptotrichia* and *Capnocytophaga* genera (**Fig. S8C**), which were similarly found to exhibit high/medium confidence enrichment in the plaque specimens using differential abundance analyses (**Fig. 4**). However, topic 1 also included *Prevotella* as a major driver, despite enrichment of multiple *Prevotella* species in the abscess (**Fig. 5**). As *Prevotella* is the second-most abundant genus in the oral cavity, it contains a wide variety of both commensal and inflammophilic species (29,82), which presumably contributed to the LDA output. We had noted a similar finding with the *Fusobacterium* genus as well (**Fig. S8C**). Our results suggest that LDA analysis is an underutilized and broadly useful metric for revealing the co-occurring polymicrobial signatures present in clinical specimen cohorts. Importantly, LDA topic modeling of microbiome data captures the strengths of standard differential abundance analysis metrics, while minimizing the negative influence of the inevitable variability that occurs with human clinical specimens.

As with our use of complementary differential abundance analytic approaches, we employed multiple independent metrics to evaluate the metabolic potential of plaque and abscess communities. Both PICRUSt2 (**Table S6**) and Anvi’o (**Table S9**) were used to analyze the distribution of KOs within the oral specimens, and the results of the enrichment analysis were further mapped onto representative KEGG pathways (**Table S7; Table S10**). Both methods have differing limitations (48) but revealed similar trends in community metabolic potential. Dental plaque communities are enriched in biosynthetic anabolic functions used to generate essential metabolites as well as energy-generating catabolic pathways targeting common dietary carbohydrates, implying a fast-growing, highly interdependent microbial ecology supported by interspecies metabolic complementation. In contrast, abscess communities exhibited a conspicuous enrichment of catabolic functions and motility pathways that are likely associated with host tissue destruction, as well as resistance mechanisms targeting host-derived antimicrobials. These enrichment patterns likely reflect the toxic inflammatory environment of the abscess and the unique nutrient conditions created as a consequence of host inflammatory responses (16,83). It remains to be determined whether the species composition of inflammophilic communities is principally attributable to analogous metabolic interdependencies like those of eubiotic communities, or if the composition of inflammophilic communities is simply a consequence of selective growth advantages for individual species that can both resist innate immunity and survive upon the nutrients provided by host inflammatory responses. For example, the specific depletion of highly abundant oral commensal species at sites of inflammation presumably relieves antagonistic growth competition with many pathobiont species. However, it may also force an obligate dependence upon inflammation and/or immunopathology to provide the critical metabolites that would otherwise have been synthesized by cohabitating commensal species. Such a result might explain the lack of enriched biosynthetic pathways in the abscess communities. It is conceivable that similar ecological principles to those described in the current study may also define the emergence of the dysbiotic microbiome communities commonly observed at other sites of inflammatory dysbiosis throughout the body. Further insights into the characteristic metabolic capacities supporting inflammophilic communities may ultimately reveal new strategies to disrupt them by depriving access to key metabolites uniquely supplied by the host.

## Data availability

All code is publicly available on github (https://github.com/kriegerm/PA_cohort_analysis). 16S rRNA V3V4 gene sequencing reads have been deposited on Sequence Read Archive in PRJNA1312071.

## Acknowledgments

Funding for this work was provided by NIDCR/NIH awards DE028252 to Justin L. Merritt, DE029612 and DE029492 to Jens Kreth, DE030859 to Madeline Krieger, and R01 DE031470 to Jeffery S. McLean. The research reported in this publication used computational infrastructure supported by the Office of Research Infrastructure Programs, Office of the Director, of the National Institutes of Health under Award Number S10OD034224. The content is solely the responsibility of the authors and does not necessarily represent the official views of the National Institutes of Health.

## Author contributions

Conceptualization M.K.; Manuscript writing M.K., G.G.Y., J.L.M.; Data generation and analysis M.K.; Figure generation M.K.; Computation M.K. and G.G.Y.; Clinical specimen collection E.A.P.; Next-generation sequencing K.A.K. and J.S.M.; Review and editing by all authors.

---

**Figure S1.**
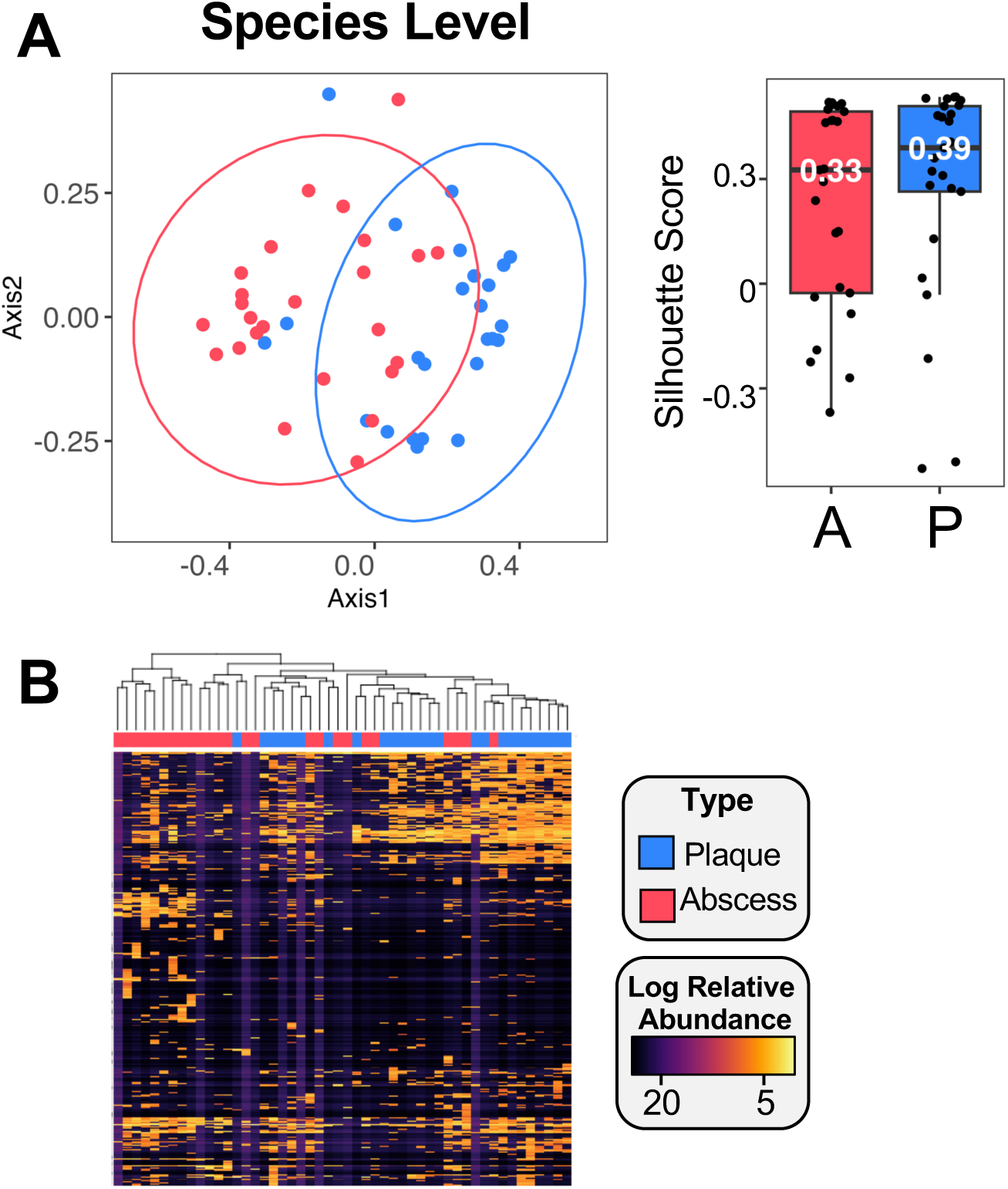
Species-level ecological composition of plaque and abscess communities. (A) Principal coordinate analysis (PCoA) of species relative abundance values using Bray–Curtis distance with silhouette scores shown alongside the PCoA plot. (B) Hierarchical clustering (Pearson correlation) of log-transformed relative abundances of species separates plaque and abscess communities.

**Figure S2.**
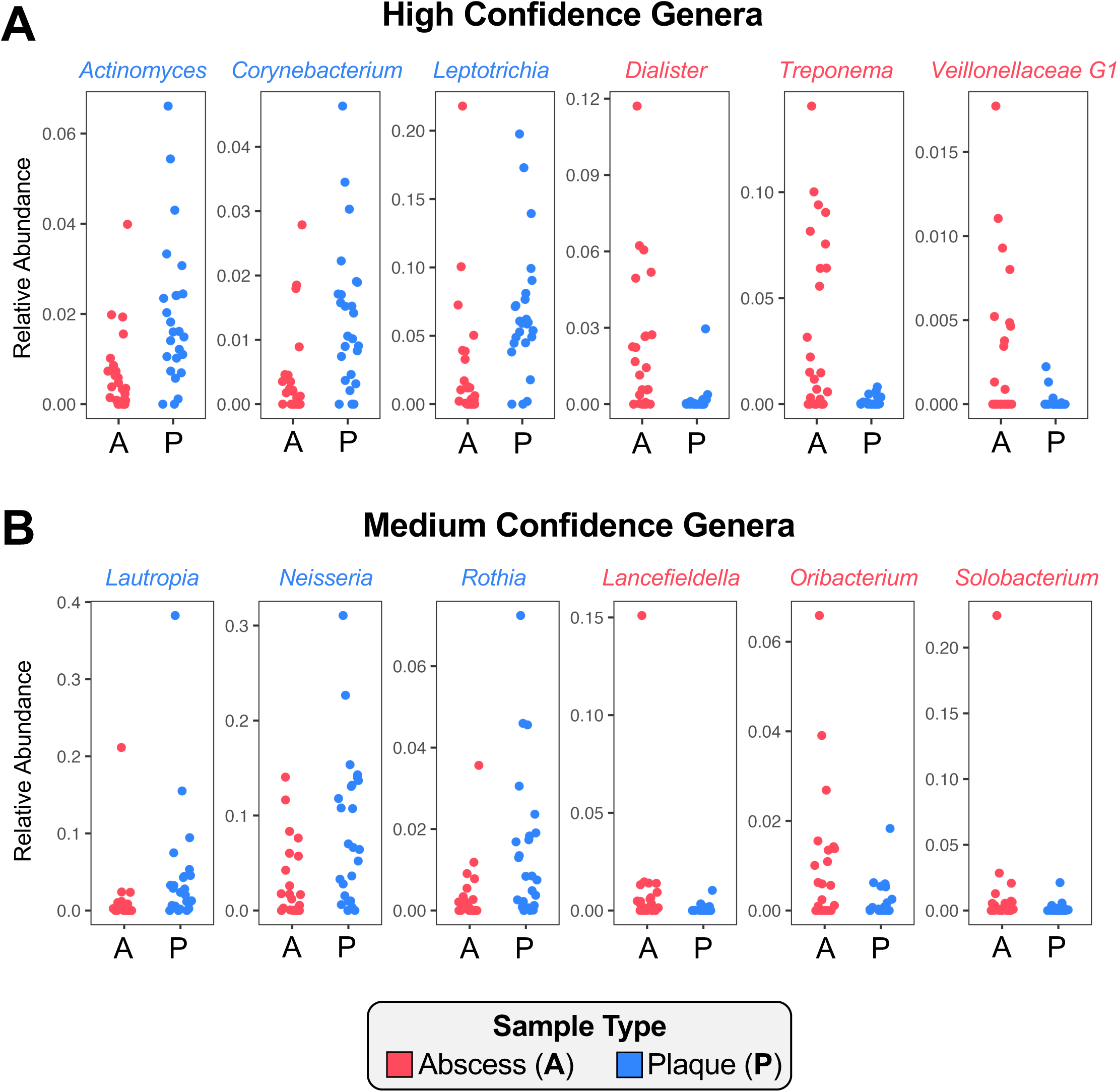
Trends in high and medium confidence differentially abundant genera. Per-sample dot plots of the relative abundance values for selected genera identified as (A) high confidence and (B) medium confidence from Figure 4 show stable trends across the entire cohort. Genera identified as being enriched in the abscess are labeled in red, while those enriched in the plaque are labeled in blue.

**Figure S3.**
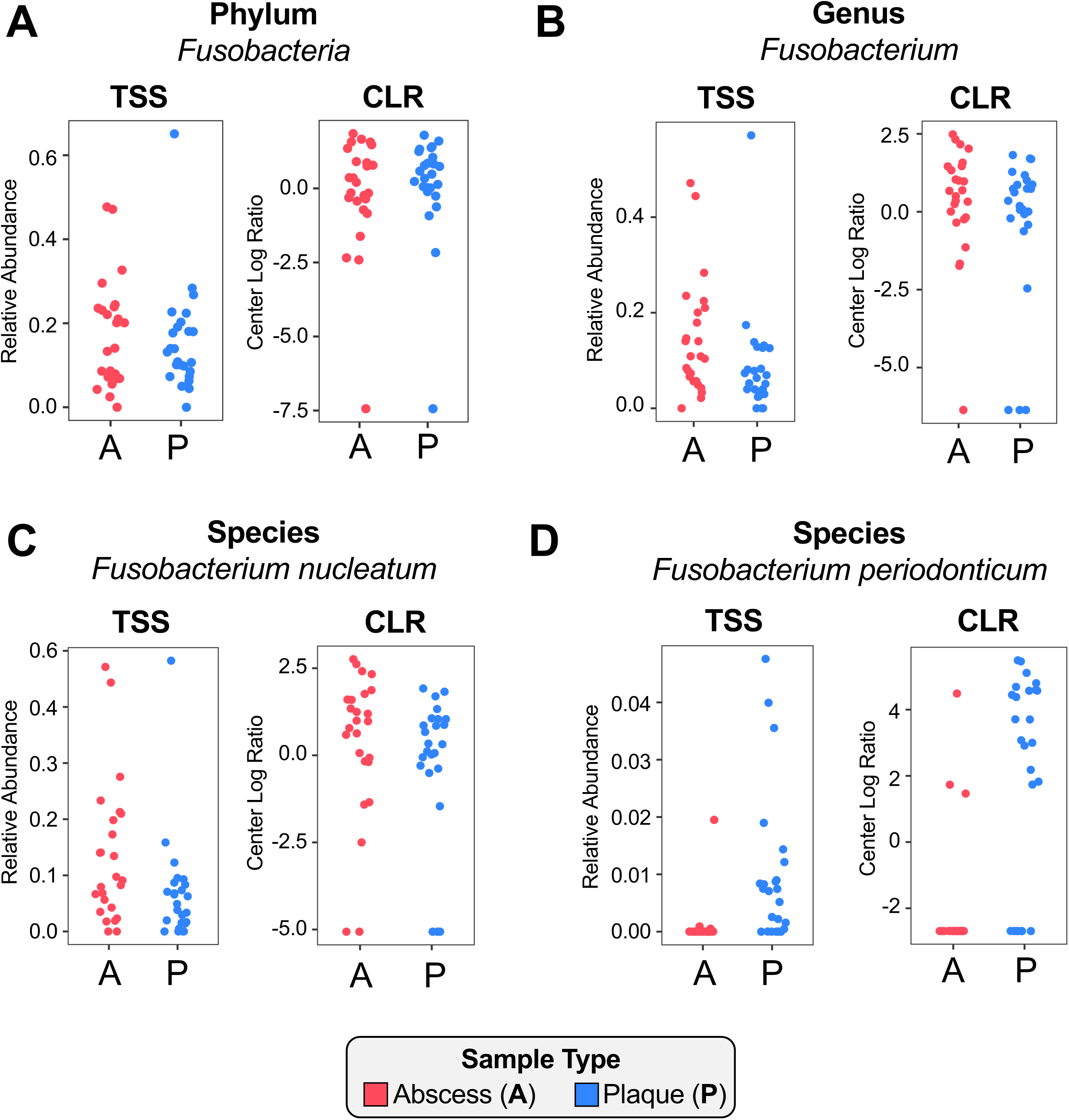
*Fusobacterium* abundance measured at different taxonomic levels. Relative abundances of fusobacteria are shown at the (A) phylum, (B) genus, and species levels for (C) *Fusobacterium nucleatum* and (D) *Fusobacterium periodonticum*. Total sum scaling (TSS) values are displayed on the left of each panel, with centered log-ratio (CLR) transformed values shown on the right.

**Figure S4.**
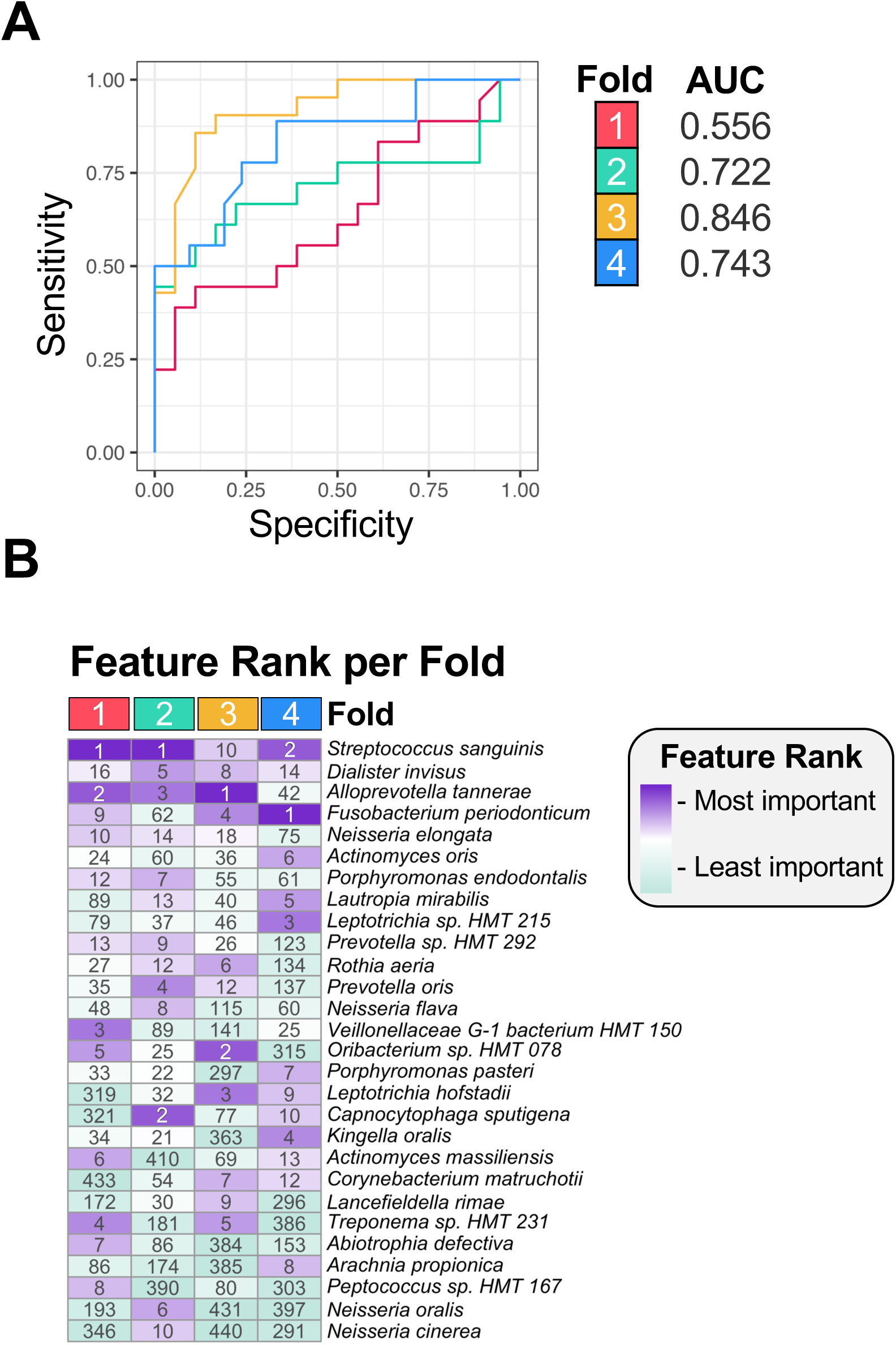
Comparison of predictive accuracy for genus- and species-level sequence data. In contrast to the random forest model built from genus-level data (Fig. 6), sequence data agglomerated at the species-level was less useful for classifying community type. (A) Receiver operating characteristic (ROC) curves from 4-fold cross-validation exhibit strong performance (area under the curve, or AUC values of 0.56–0.85). (C) Heatmap of feature importance rankings across folds, with top predictors in purple and lower-ranked features in green. The rank of feature importance is shown inside each cell.

**Figure S5.**
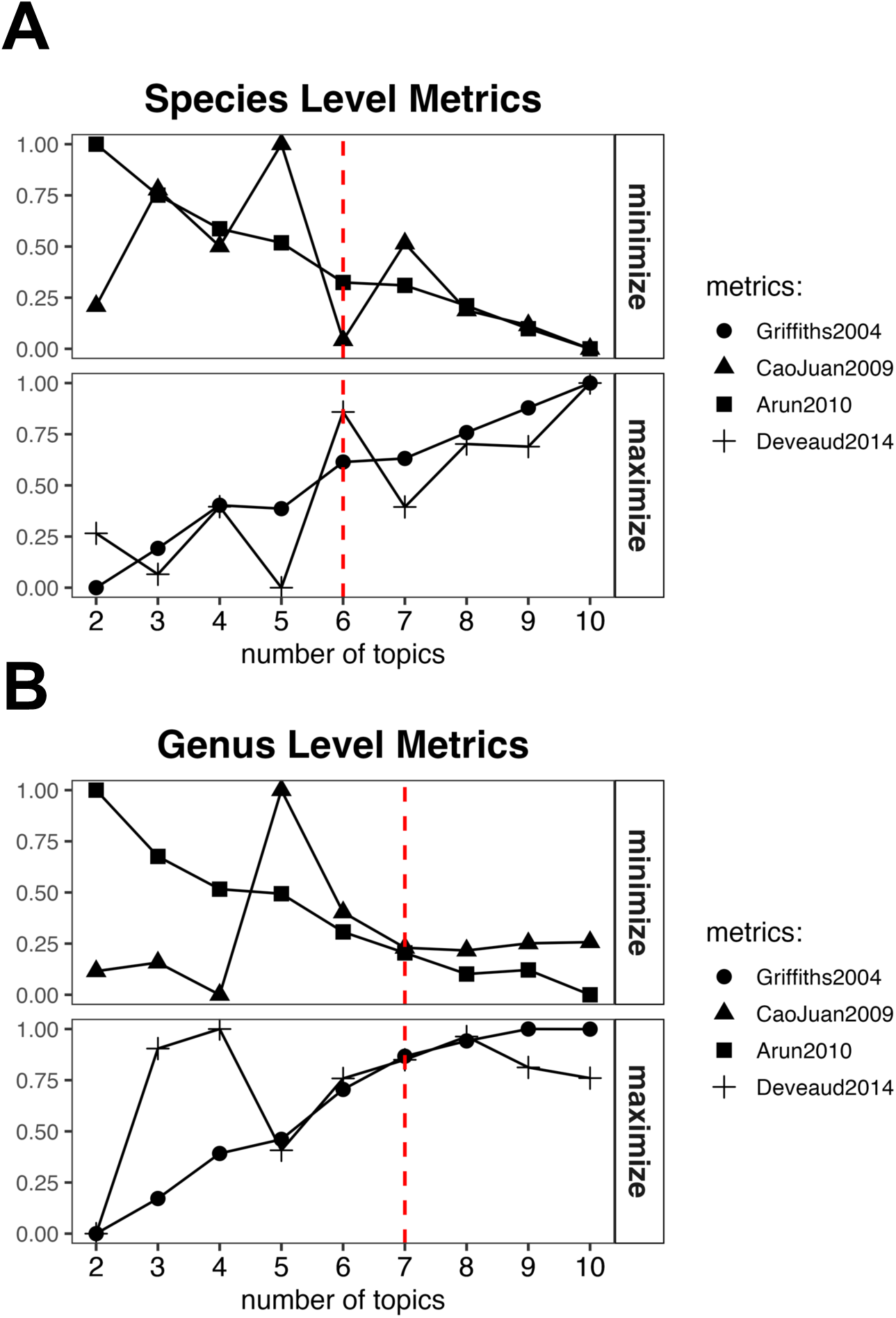
Algorithmic validation metrics used to select the optimum number of topics for Latent Dirichlet Allocation (LDA) topic-modeling. Four different metrics were used to validate the performance of an LDA model used for (A) species-level and (B) genus-level data. To preserve interoperability of the model, the number of topics minimized CaoJuan2009 (84) and Arun2010 (85) metrics and maximized Deveaud2014 (86) and Griffiths2004 (87) metrics. The number of topics selected is shown with a red dotted vertical line.

**Figure S6.**
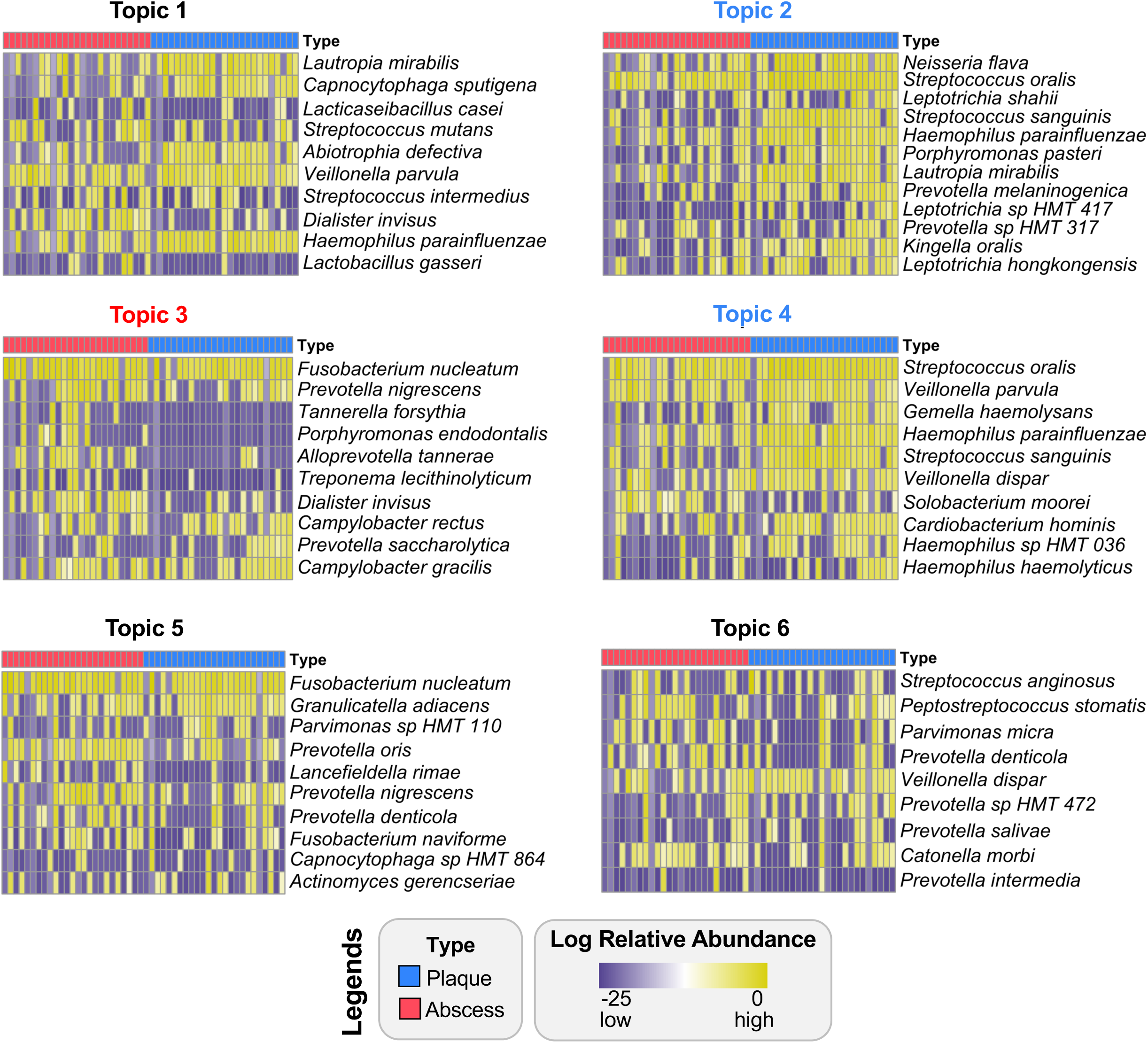
Patterns of species-level enrichment across six topics. Relative abundance log values of the top taxa identified as significant drivers of each topic signature. Topics 2 and 4 (blue text) are significantly enriched in plaque specimens and topic 3 (red text) is significantly enriched in abscess communities.

**Figure S7.**
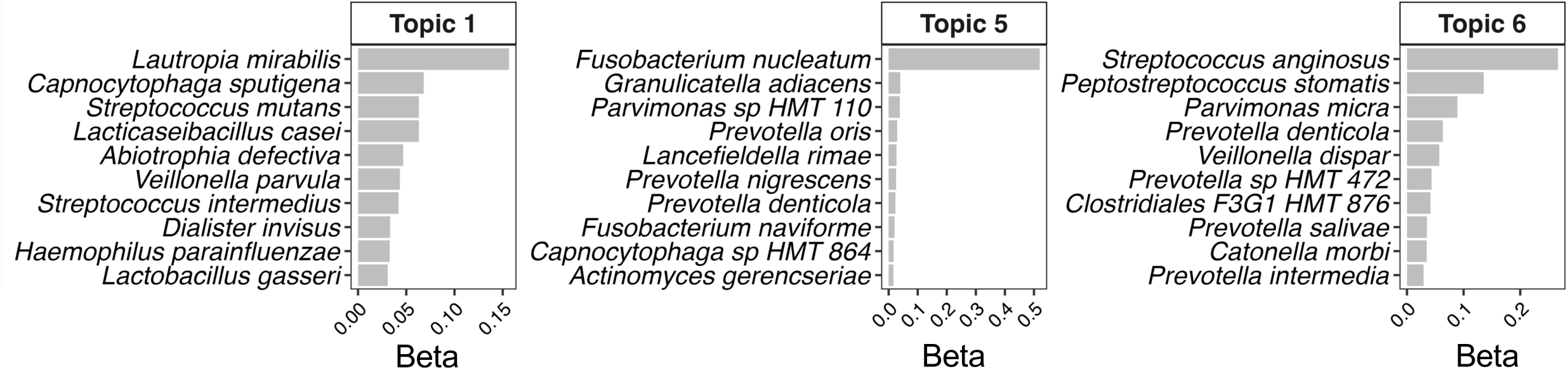
Importance scores of species-level topics that are shared equally between plaque and abscess specimens. The importance (beta) scores of the top species driving topics 1, 5 and 6 are shown. These topics were not significantly enriched in either plaque or abscess communities but instead describe shared signatures of both oral communities.

**Figure S8.**
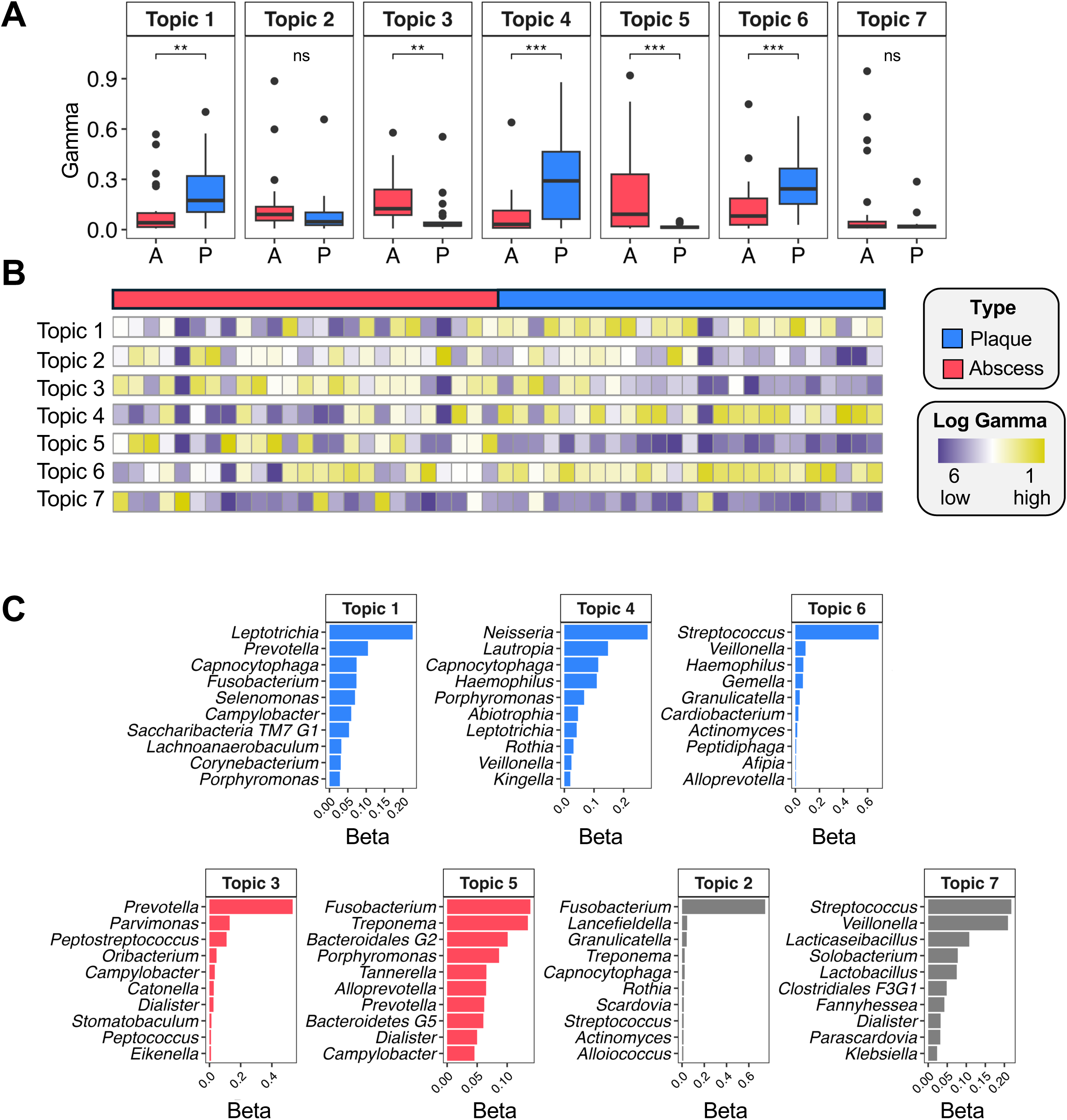
Topic modeling of genus-level community signatures in plaque and abscess specimens. (A) Boxplots of sample membership (gamma values), with the results of a Wilcoxon Signed-rank test displayed above (*** = p < .001, * = p < .05). (B) Heatmap of gamma scores across topics. (C) The importance (beta) scores of the top species driving each topic. Topics 1, 4 and 6 were enriched in plaque samples, while topics 3 and 5 were enriched in abscess communities. Topics 2 and 7 described a shared microbiome signature across both plaque and abscess specimens.

**Figure S9.**
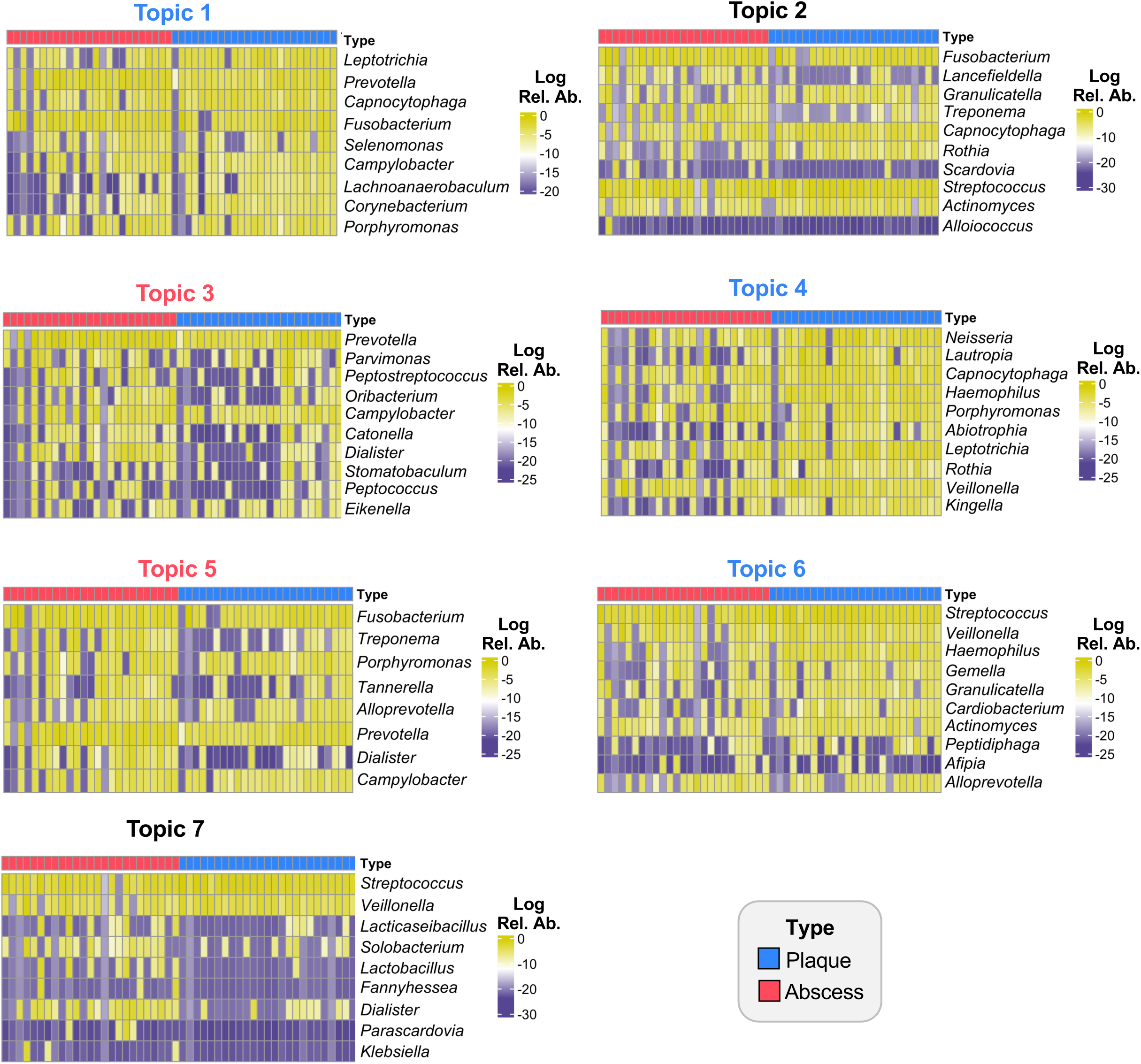
Patterns of genus-level taxa enrichment across seven topics. Log values of the relative abundances of the top taxa identified as significant drivers of each topic signature. Topics 1, 4 and 6 (blue text) are significantly enriched across plaque specimens and topics 3 and 5 (red text) are significantly enriched in abscess specimens.

**Figure S10.**
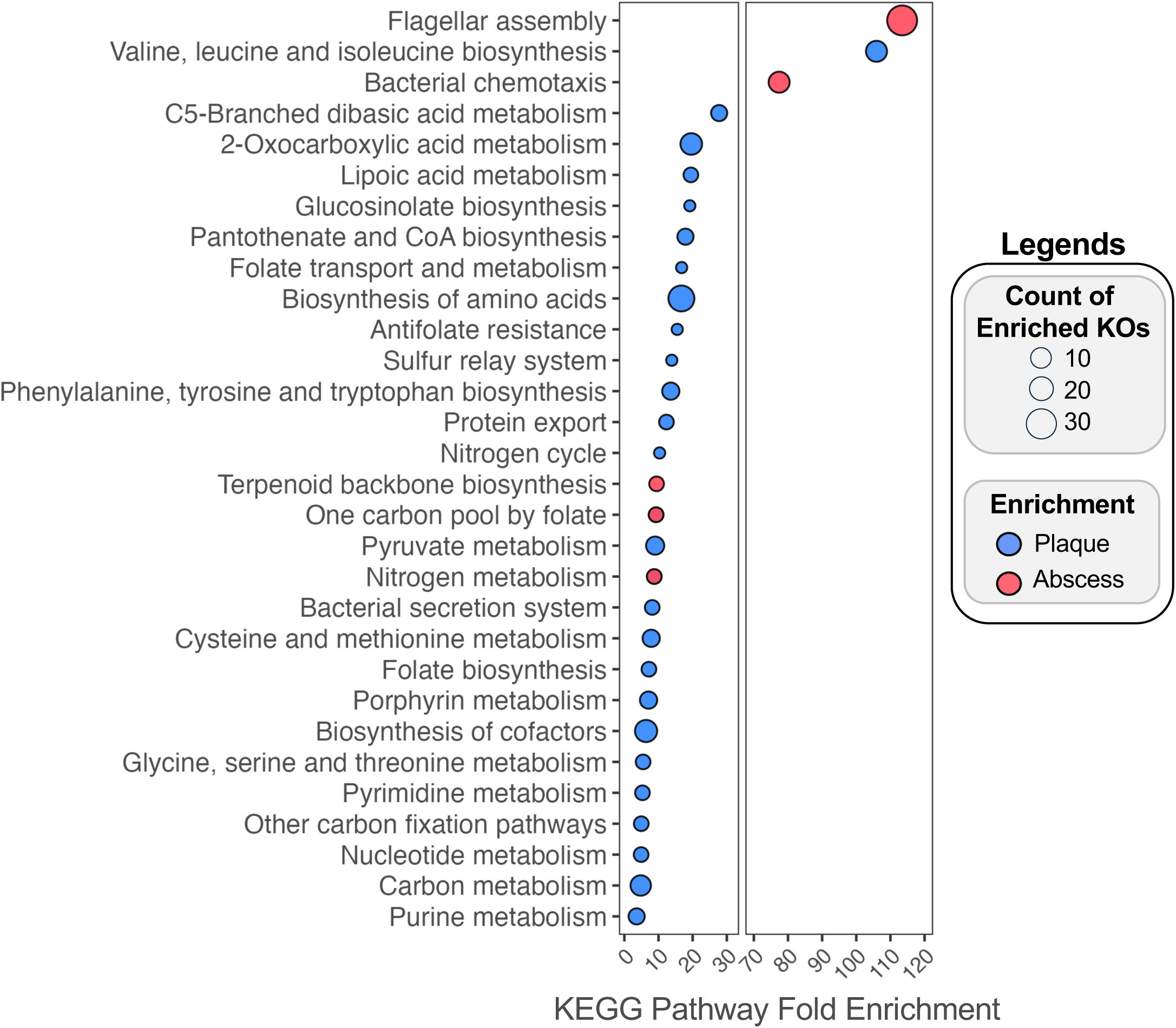
Additional confirmation of KEGG pathways within plaque and abscess communities. A functional enrichment analysis was performed on a pangenome constructed of enriched taxa identified in the plaque and abscess specimens. KOs that were differentially enriched either plaque and abscess samples (adjusted q value < .05) were used as inputs to identify key KEGG pathways of plaque (blue) and abscess (red) communities. The x-axis displays fold enrichment of the KEGG pathway. Dot size corresponds to the number of KOs that were identified as being part of each respective KEGG pathway (y-axis).

## Tables

**Table S1. Genus-level differences in mean and median values between plaque and abscess communities.**

**Table S2. Counts of patients having enrichment of taxa in either the plaque or abscess specimens at the genus- and species-levels.**

**Table S3. Species-level differences in mean and median values between plaque and abscess communities.**

**Table S4. Differential abundance results from three analytical methods.**

**Table S5. Counts of patients harboring enriched taxa identified by differential abundance analysis of plaque and abscess specimens.**

**Table S6. MaAsLin2 analysis of KEGG Orthologs identified by PiCRUSt2.**

**Table S7. Enriched KEGG Pathways in plaque and abscess samples identified by PiCRUSt2.**

**Table S8. Genomes used for Anvi’o functional enrichment analysis.**

**Table S9. Complete results from Anvi’o functional enrichment analysis of KEGG Orthologs.**

**Table S10. Enriched KEGG Pathways in plaque and abscess samples identified by Anvi’o.**

**Table S11. Contaminant taxa removed from analysis.**

## Materials and Methods

### Sample collection

As part of standard clinical care for one or more abscessed teeth, deidentified clinical specimens were collected from pediatric dental patients at Oregon Health & Science University (OHSU) dental clinics. Prior to the start of the project, all protocols were reviewed by the OHSU IRB and classified as “Not Human Subjects Research”. Supragingival plaque was collected with a sterile swab from the facial surface of a non-carious primary tooth and abscess samples were collected from the furcation of an extracted primary tooth. In the event of a draining abscess, purulence was sampled from the site of the fistula. Samples were transferred to the anaerobic chamber, mixed with glycerol freezing media, and stored at −80 °C.

### DNA collection and sequencing preparation

DNA was isolated from samples stored in DNA/RNA shield using the ZymoBIOMICS DNA Miniprep kit (Zymo Research) per the manufacture’s recommendation, using a double-elution protocol with 50 µl of H2O warmed to 60°C and no Zymo-Spin III-HRC filtering step. 16S rRNA gene sequencing was performed as previously reported (88,89). Briefly, the V3-V4 variable region of the 16S rRNA gene was amplified using gene-specific primers with Illumina adapter overhang sequences

(5′-TCGTCGGCAGCGTCAGATGTGTATAAGAGACAGCCTACGGGNGGCWGCAG-3′ and 5′-GTCTCGTGGGCTCGGAGATGTGTATAAGAGACAGGACTACHVGGGTATCTAATCC-3′) using either 12.5 µl Kapa HiFi Hot Start ReadyMix, 2.5 µl gDNA, and 10 µl sterile H_2_O or with the following protocol: 95°C for 3 min; either 35 or 40 cycles at 95°C for 30 seconds, 55°C for 30 seconds, 72°C for 30 seconds; 72°C for 5 min using 40 cycles of amplification due to low bacterial abundance in the original sample. Amplicons were cleaned using AMPure XP beads and eluted into 50 µl of H_2_O. Indices were attached using 8 cycles of the same PCR protocol, and amplicons were cleaned with AMPure XP beads. The size of select products were verified using gel electrophoresis. 25 µl of each sample was submitted to the McLean Lab at the University of Washington for Illumina MiSeq sequencing (2×300bp, 10% PhiX spike).

### Community analysis and bioinformatics

QIIME2 (version 2022.2.0) (90) and DADA2 (91) within the QIIME2 package were used to trim and classify sequences using the feature-classifier classify-consensus-blast referencing the Human Oral Microbiome Database (version 15.23). R studio Phyloseq (version 1.52.0) (92) was used for subsequent analysis. Contaminates listed in **Table S11** were removed from the analysis, and ASVs found only in the PCR and reagent blank samples were removed. Additionally, only samples where both plaque and abscess samples contained >1,000 reads were retained. The ggplot2 and pheatmap packages in R were used to produce figures.

### Differential abundance analysis

Three differential abundance algorithms were tested: Microbiome Multivariable Associations with Linear Models (MaAsLin2, version 1.22.0) (35), Analysis of Compositions of Microbiomes with Bias Correction (ANCOMBC2, version 2.10.2) (36), and Analysis Of Differential Abundance Taking Sample and Scale Variation Into Account (ALDEx2, run through the microbiomeMarker package version 1.13.2) (37). Model parameters for MaAsLin2 and ALDEx2 were chosen through parameter sweeps, which considered the number of differential taxa found through each combination of data transformation, normalization, and method. MaAsLin2 was run with a linear model, CSS normalization, and log transformation of count data with a q-value significance threshold of 0.05. ALDEx2 was run using a log10 transformation, CSS normalization of count data, and Wilcoxon Signed-rank test to assess significance with a Benjamin-Hochberg adjusted p-value threshold of 0.05. ANCOMBC2 used a Holm p-value adjustment method and a 0.05 significance threshold.

### Ordination

To choose the ordination method and parameters, we used an iterative parameter sweep to select the most appropriate distance measure and data transformation for each ordination method (Principal Coordinates Analysis, Principal Component Analysis, or Non-metric multidimensional scaling). This parameter sweep found the combination of distance measures (Jaccard, Bray-curtis, or Euclidean) and data transformations (total sum scaling and center log ratio) that minimized the correlation (spearman’s rho) with library size for the first axis of variation, maximized the silhouette score within each type (plaque and abscess), and considered the percent variation captured by the first two axes. Considering these parameters, we chose PCoA of relative abundance values using bray-curtis distance as our final ordination model.

### Random Forest

Relative abundance taxa values were used as input to the random forest model agglomerated at the genus level. The caret package in R (version 7.0-1) was used to construct a random forest model using 4-fold cross validation predicting the plaque or abscess status of each sample.

### Metabolic Profiling

PICRUSt2 (48) (version 2.6.2) was run from a conda environment using trimmed representative sequences output by the Qiime2 pipeline. The results of the KO analysis were analyzed for differential abundance using MaAsLin2 (CLR normalization, no transformation). Lists of significant results (q value < .05) for KOs differentially enriched in plaque and abscess samples (**Table S6**) were used for input into KEGG enrichment analysis using the ClusterProfiler enrichKEGG command. Results for both the plaque and abscess specific analysis were combined and shown in Figure 8, with the fold enrichment of these KEGG pathways on the x-axis.

An Anvi’o (62) (version 8) pangenome was constructed from all the species identified as being differentially abundant in either plaque or abscess communities using any of the three methods (**Table S4**). RefSeq representative genomes for each species were collected from NCBI (**Table S8**). Taxa without any available genomes were omitted from the pangenome analysis. Anvi’o was used to construct the pangenome, and functional enrichment in either plaque or abscess communities was computed for KOs using the anvi-compute-functional-enrichment-in-pan (64) command. Functional enrichment results were filtered by adjusted q-value (q < .05) (**Table S10**). As with the Picrust2 results, KOs that were differentially enriched in plaque and abscess samples were used as input into the enrichKEGG function from ClusterProfiler, and the resulting enrichments are shown in **Figure S10.**

### Topic Modeling

For taxa, relative abundance values agglomerated at the genus level were used as input for topic modeling. A scaling factor of 1,000 was chosen based on silhouette score optimization for both plaque and abscess communities. The number of topics was determined from a set of four validation metrics (84–87). The LDA function from the topicmodels package (version 0.2-17) was used to build an LDA models for species and genus level data. A Wilcoxon Signed-rank test with FDR p-value correction was used to assess the differential significance of topic membership by type.

